# Natural aging drives a subclinical cardiovascular phenotype in a non-human primate

**DOI:** 10.64898/2026.05.08.723137

**Authors:** Lina Klösener, Mostafa Samak, Dominik Lerm, Jia Li Ye, Federico Bleckwedel, Amir Moussavi, Tor Rasmus Memhave, Majid Ramedani, Fernanda Ramos Gomes, Amara Khan, Ajinkya Kulkarni, Maren Sitte, Gabriela Salinas, Jessica König, Wiebke Möbius, Boguslawa Sadowski, Sabine Steffens, Meik Kunz, Laura Zelarayan, Christof Lenz, Christian Bär, Rüdiger Behr, Susann Boretius, Frauke Alves, Thomas Thum, Giulia Germena, Matthias Mietsch, Rabea Hinkel

## Abstract

Aging is an inevitable risk factor for cardiovascular disease. Profound understanding of mechanisms underlying the early stages of cardiovascular aging is essential for the development of novel therapeutics. Therefore, animal models which closely reflect the human condition are highly sought after. Here, we investigated natural cardiovascular aging in a non-human primate, comparing healthy young-adult and aged common marmosets (*Callithrix jacchus*). Despite preservation of most cardiac functional parameters in aged animals, significant histological alterations were found including fibrosis and microvascular rarefaction. Molecular phenotyping by single-nuclei RNA-sequencing revealed activation of cardiac stress, pro-inflammatory and fibrotic gene programs in aged hearts. Importantly, proteomic analysis of cardiac extracellular vesicles revealed a cardioprotective cargo in young animals while functionally demonstrating pro-angiogenic properties on human cardiac microvascular endothelial cells. Finally, large vessel atherosclerosis was strikingly evident in aged animals and elucidated by bulk RNA-sequencing. Overall, the aging marmoset offers a great potential for translational cardiovascular research.

## Introduction

Humanity has made remarkable feats aiming to extend and improve the quality of life. Yet, cardiovascular diseases linger as the leading cause of morbidity and mortality worldwide, with aging being an inevitable risk factor influencing their prevalence and severity^1–3^. A mounting body of knowledge has accumulated over decades of biomedical research addressing several cardiovascular disease entities and their comorbidities. However, little is known about natural aging in and of itself as an independent driver of cardiovascular pathology in the absence of other risk factors. To study such gradual and complex process as cardiovascular aging, animal models that closely mirror the human situation are sought after.

The common marmoset (*Callithrix jacchus*) is a New World monkey, which combines the advantages of research in non-human primates (NHP), being humans’ closest evolutionary relatives, with those of non-primate species. These include a shorter life span, easier husbandry and handling, as well as lower costs compared to other larger primates. Indeed, recent studies demonstrated similarities between common marmosets and humans in cardiac anatomy and topography, as well as age-related functional changes^4–9^. Moreover, the possibility to generate genetically modified marmosets further expands their research application potential^10,11^. Therefore, characterization of the cardiovascular aging phenotype in this model can be of great translational utility, guiding promising therapeutic strategies in humans.

Here, we took a comprehensive approach to study aging of the cardiovascular system in marmosets, combining *in vivo* cardiac functional analysis and histology with single-nuclei RNA-sequencing (snRNA-seq). In addition, we addressed cardiac intercellular communication through proteomic analysis of cardiac extracellular vesicles (EVs), emphasizing differential protein cargo between age groups, and demonstrating an *in vitro* effect on primary human cardiac microvascular endothelial cells (HCMEC). Finally, we characterized age-driven vascular pathology by thorough histological analysis combined with bulk RNA-sequencing of thoracic and abdominal aortas.

## Results

### Cardiac functional and blood parameters of healthy aged common marmosets reveal a mild clinical phenotype

In the present study, healthy young-adult (< 4 years) and aged (> 11 years) common marmosets were included (Fig. 1a). The animals underwent clinical and hematological examinations to confirm their health status (Fig. 1b–h, Supplementary Fig. 1a–g). Fasted blood parameters showed no differences in C reactive protein (CRP), leukocyte count, liver enzymes or kidney function (Fig. 1b–g)^12^. High density lipoprotein (HDL), however, was significantly lower in aged animals, resulting in an elevated non-HDL/HDL ratio, pointing towards altered cholesterol metabolism (Fig. 1h, Supplementary Fig. 1h–l).

**Figure 1:**
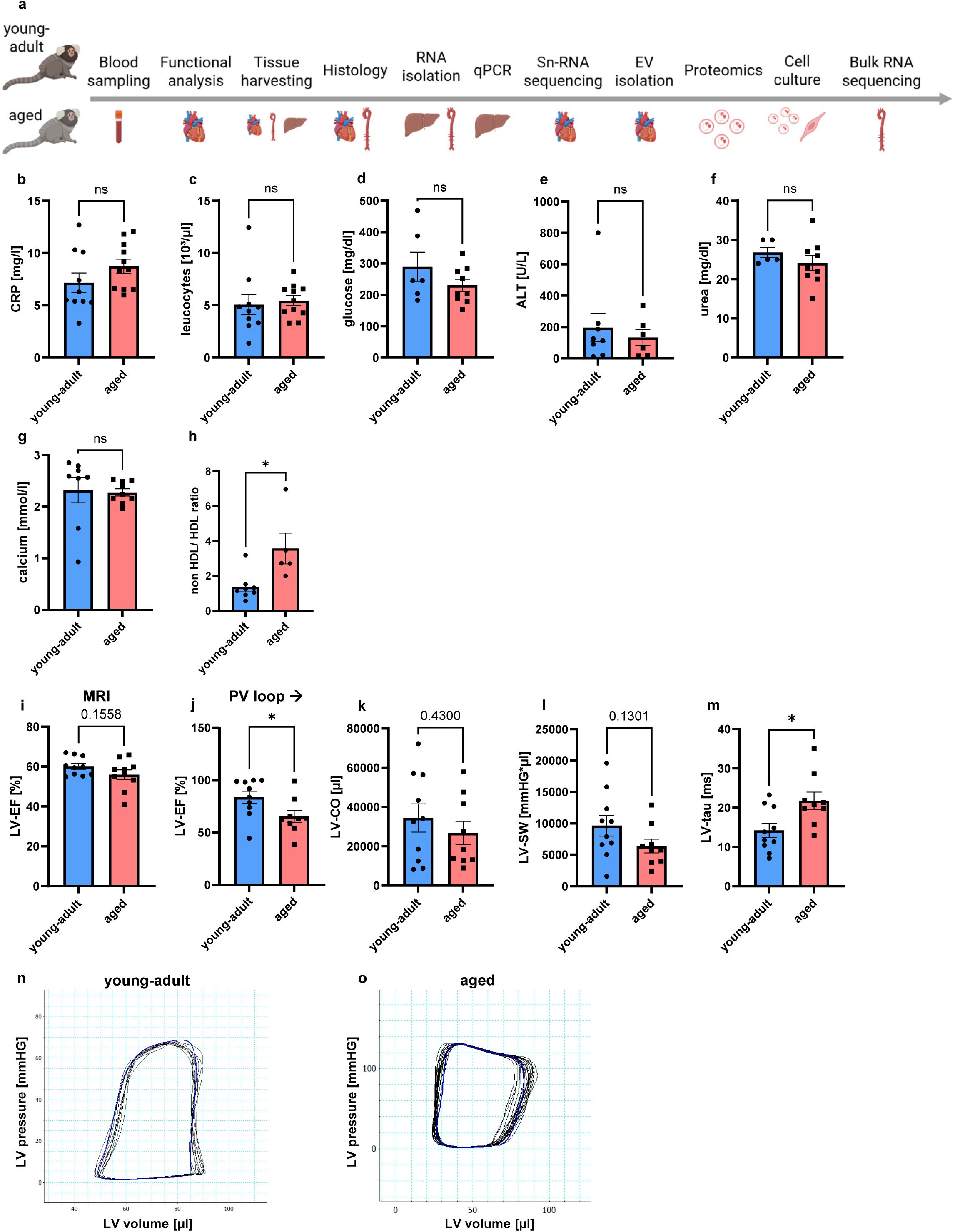
Study design, blood analysis and assessment of cardiac functional parameters. **a**, Study design (illustrated in BioRender). **b-h**, Blood analysis of C-reactive protein (CRP; young-adult n = 10, aged n = 11) (**b**), leucocyte count (young-adult n = 10, aged n = 11) (**c**), glucose (young-adult n = 6, aged n = 9) (**d**), alanine aminotransferase (ALT; young-adult n = 8, aged n = 6) (**e**), urea (young-adult n = 5, aged n = 9) (**f**), calcium (young-adult n = 8, aged n = 9) (**g**) and non-HDL/HDL ratio (young-adult n = 8, aged n = 5) (**h**). **i**, Cardiac MRI assessment of left-ventricular ejection fraction (LV-EF; n = 10). **j-m**, Intra-cardiac pressure-volume loop (PV loop) measurments (young-adult n = 10, aged n = 9) of LV-EF (**j**), cardiac output (LV-CO; **k**), and stroke work (LV-SW; l) and the time constant of isovolumic relaxation (LV-tau) (**m**). Bar graphs represent means ± SEM. Statistical significance was assessed by unpaired twotailed Student’s t-tests (\**P* < 0.05, ns: not significant). **n,o**, Representative PV loops from one young-adult and one aged animal.

Cardiac function was investigated using magnetic resonance imaging (MRI) and invasive intra-cardiac pressure-volume loop (PV loop) catheter. Neither showed differences in left ventricular ejection fraction (LV-EF), cardiac output (LV-CO) nor stroke work (LV-SW) (Fig. 1i,k,l), whereas PV loop revealed a significant decrease of EF in aged animals (65% in aged vs. 83.7% in young animals, *P* < 0.05) (Fig. 1j). Moreover, aged animals displayed prolonged Tau (LV-Tau), the isovolumic relaxation constant, upon PV loop measurement, indicating impaired diastole (Fig. 1m–o, Supplementary Fig. 2a–d).

### Age-driven cardiovascular morphological changes in the common marmoset

Assessment of cardiac morphological changes revealed significant changes between healthy young-adult and aged animals (Fig. 2,3). MRI of extracellular volume (ECV), a marker for fibrosis, revealed increased values for aged animals (Fig. 2a,b). To confirm cardiac fibrosis in aged marmosets, whole heart cross-sections were analyzed for collagen content via Masson trichrome staining showing significantly larger collagen areas compared to young animals (Fig. 2c,d)^13^. Results from both atria and ventricles corroborated the findings from whole-heart cross-sections (Fig. 2e–l). Additionally, perivascular fibrosis was evident in aged animals’ left ventricles, as revealed by higher perivascular collagen content (Fig. 2m,n, Supplementary Fig. 2e). Cardiac fibrosis can impair myofibril organization^14^. To characterize such structural alterations, we performed label-free two photon microscopy followed by alignment analysis. Values close to 1 indicate oriented alignment, whereas wavier structures assume lower values, indicative of poor alignment^15^. Indeed, aged animals displayed wavier cardiomyocyte myofibrils compared to well-aligned myofibrils in young hearts, corroborating the connection between cardiac fibrosis and myofibril disorganization (Fig. 2o,p).

**Figure 2:**
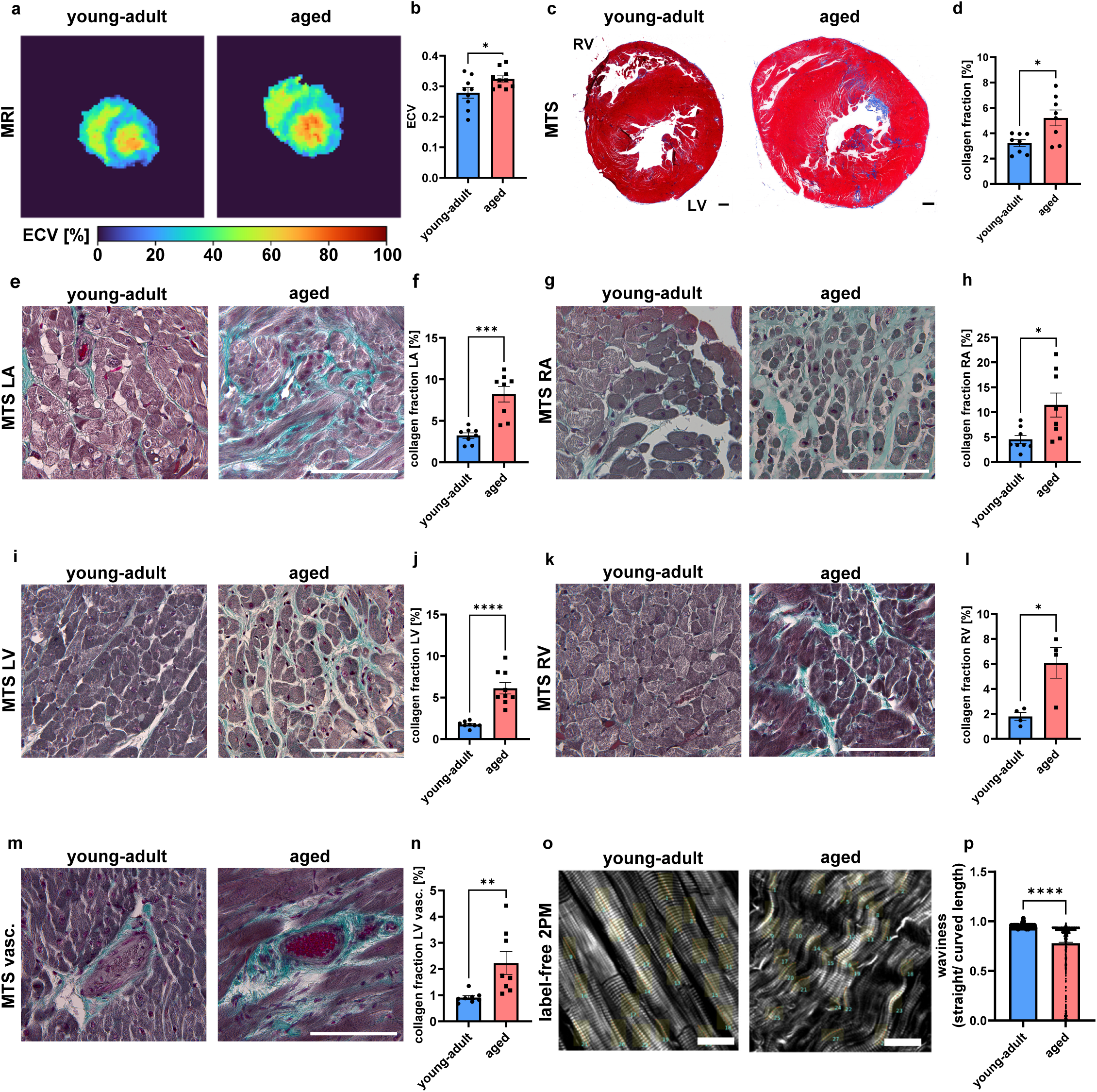
Cardiac histological assessment of extracellular matrix deposition and cardiomyocyte structure. **a,b**, MRI assessment of extracellular volume (ECV) (young-adult n = 9, aged n = 10). **c**,**d**, Masson trichrome staining (MTS) (**c**) and collagen fraction quantification (n = 8) (**d**) in whole heart cross-sections shortly below valve level. Scale bar = 1 mm. **e**-**n**, MTS and collagen fraction quantification in left atria (**e**,**f**) and right atria (n = 8) (**g**,**h**), left ventricles (young-adult n = 8, aged n = 9) (**i**,**j**) and right ventricles (n = 4) (**k**,**l**), and perivascular region (n = 8) (**m**,**n**). Scale bar = 100 μm. **o**,**p**, Label-free two-photon microscopy (2PM) imaging of LV (**o**), and waviness analysis of traced myofibrils (young-adult n = 1331, aged n = 849) (**p**). Scale bar = 30 μm. LA/ RA: left/ right atrium; LV/ RV: left/right ventricle. Bar graphs represent means ± SEM. Statistical significance was assessed by unpaired two-tailed Student’s t-test (\**P* < 0.05, \*\**P* < 0.01, \*\*\**P* < 0.001, \*\*\*\**P* < 0.0001).

**Figure 3:**
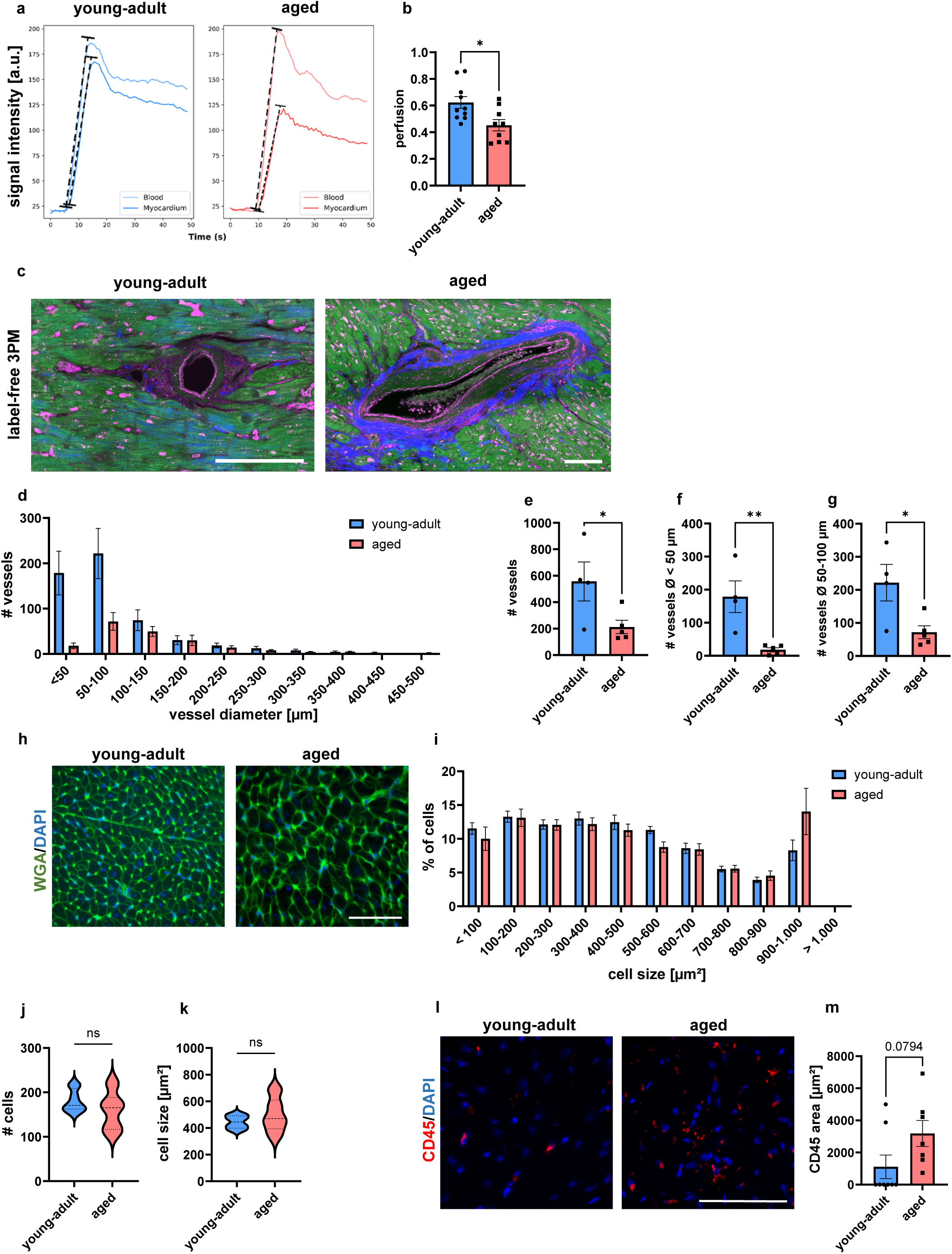
Investigation of cardiac microcirculation, hypertrophy and inflammatory state. **a-b**, Perfusion analysis by cardiac MRI (young-adult n = 10, aged n = 9). **c**, 3 Photon label-free imaging of left ventricle (LV) vascular supply (green: cardiomyocytes, blue: collagen, pink: elastin). Scale bar = 50 μm. **d**-**g**, analysis of vessel size distribution (n = 5) (**d**), total vessel count (**e**) and microvessels count with diameter < 50 μm (**f**) and diameter 50–100 μm (**g**) (young-adult n = 4, aged = 5). **h**-**k**, Cardiomyocyte size analysis by wheat germ agglutinin staining of cell membrane (green) and DAPI nuclear counterstaining (blue). Scale bar = 100 μm. LV cell size distribution analysis (**i**), cell number (**j**) and average cell size (**k**). **l**,**m**, Immunofluorescence staining of LV tissue sections for CD45 (leukocytes; red), with a nuclear counterstaining for DAPI (blue) (scale bar = 100 μm) (**l**), and analysis of CD45 area per image (young-adult n = 8, aged n = 9) (**m**). Bar graphs in **b**, **e**–**g** and **m** represent means ± SEM. Statistical significance was assessed by unpaired two-tailed Student’s t-tests (\**P* < 0.05, \*\**P* < 0.01).

Interestingly, MRI showed reduced cardiac perfusion in aged animals (Fig. 3a,b). Typically, microvascular supply is the first entity to be affected in the diseased heart, leading to reduced perfusion^16^. Herein, label-free imaging of small vessels by three-photon microscopy (Fig. 3c,d, Supplementary Fig. 2f) revealed not only a lower total number of vessels, but also a significantly reduced proportion of microvessels (≤50 and 50 – 100 µm diameter) in aged hearts (Fig. 3e–g). Microvessels with a diameter < 50 µm, while abundant in young ventricles, were scarcely detected in the aged (Fig. 3c).

As cardiac fibrosis stiffens the ventricles, cardiomyocytes adapt to the increased workload, ultimately leading to their hypertrophy. To investigate cardiomyocyte hypertrophy, heart slices were stained for cell membranes with wheat germ agglutinin (Fig. 3h). While no significant differences in average cardiomyocyte cross-sectional area were detected, there were greater variations in absolute cell numbers and sizes in older animals (Fig. 3i–k). Such heterogeneity may reflect age-related hypertrophy, where some cardiomyocytes enlarge, while others become atrophic.

Finally, leukocyte staining revealed a tendency towards inflammatory infiltration in aged hearts (Fig. 3l–m).

### Age-related atherosclerosis in the absence of comorbidities

The finding of microvascular impairment in aged hearts prompted an investigation of extra-cardiac blood vessels. Therefore, we examined thoracic aortas of young-adult and aged marmosets by histological staining to assess atherosclerotic burden (Fig. 4; Supplementary Fig. 3)^17^. Imaging revealed a presence of fatty deposits of varying extents in all aged animals, while they were absent in young animals (Fig. 4a; Supplementary Fig. 3a). Staining-based calculation of elastin/collagen ratio revealed a significant decrease in aged animals, indicative of aortic wall stiffening (Fig. 4b,c and 4i)^18^.

**Figure 4:**
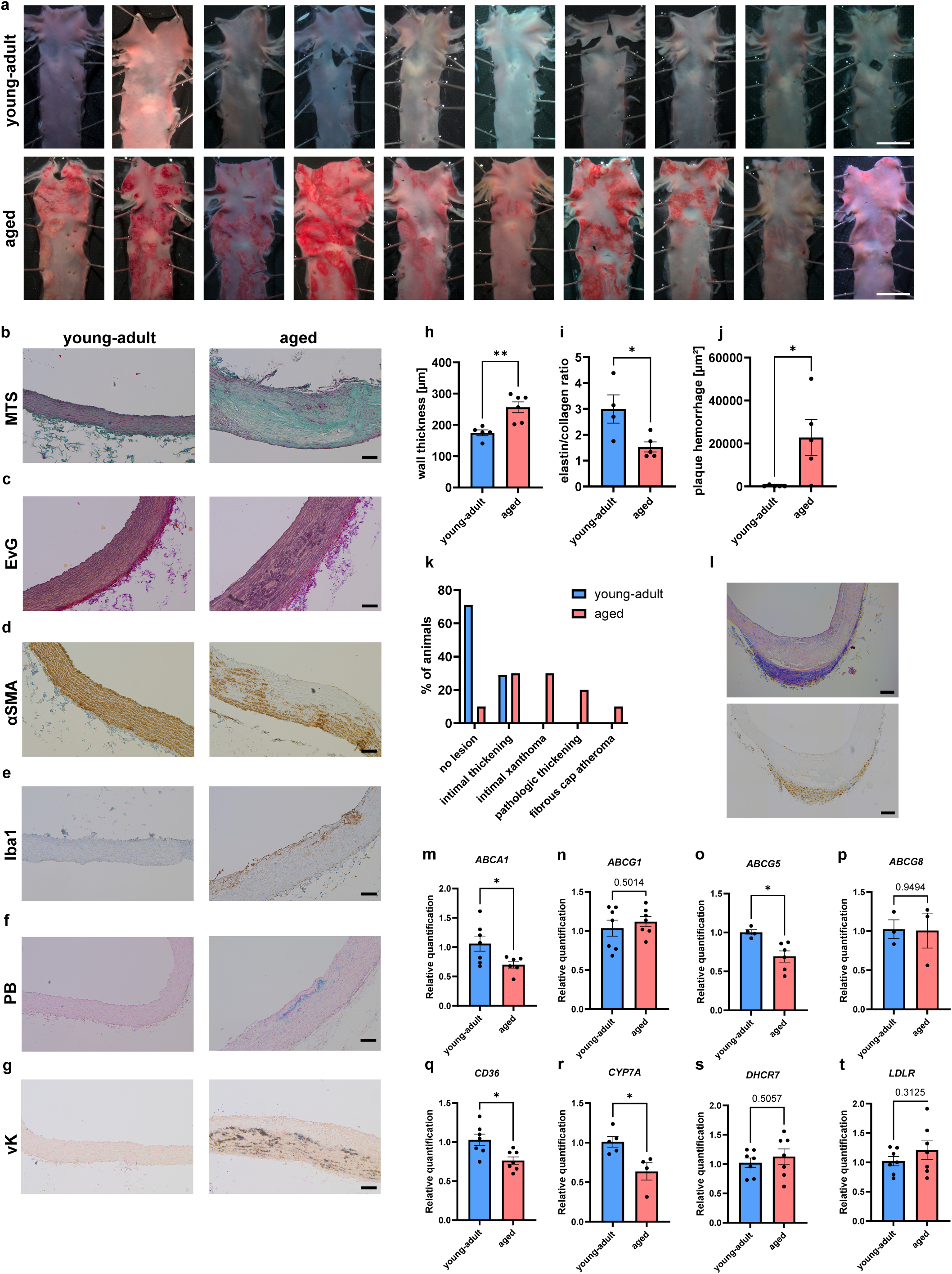
Examination of thoracic aorta and expression of hepatic cholesterol metabolic genes in young-adult and aged animals. **a**, En-face visualization of Oil Red O-stained thoracic aorta (n = 10). Scale bar = 5 mm, **b**, Masson Trichrome staining (MTS) (collagen fibres). **c**, Elastica van Gieson (EvG) staining (elastin fibers). **d**,**e**, immunohistochemistry for alpha-smooth muscle actin (αSMA) (**d**) and ionized calcium-binding adapter molecule 1 (Iba1) (macrophages) (**e**). **f**, prussian blue (PB) staining (intraplaque hemorrhage). **g**, von Kossa (vK) staining (calcification). Scale bar = 100 μm. **h**-**j**, Quantificatio of wall thickness (young-adult n = 5, aged n = 6) (**h**), elastin/collagen ratio (young-adult n = 4, aged n = 5) (**i**) and intraplaque hemorrhage (n = 5) (**j**). **k**, Plaque grading of sections according to R. Virmani (young-adult: sections n = 21; animals n = 7, aged: sections n = 29; animals n = 10). **l**, H&E staining (upper) for arterial tertiary lymphoid organs and CD3 immunohistochemistry (T-lymphocytes, lower) in an aged animal. Scale bar = 200 μm. **m**-**t**, Liver tissue qPCR for *ABCA1*, *ABCG1*, *ABCG5*, *ABCG8*, *CD36*, *CYP7A*, *DHCR7* and *LDLR* (n = 7). Bar graphs in **h**–**j** and **m**–**t** represent means ± SEM. Statistical significance was assessed by unpaired two-tailed Student’s t-tests (\**P* < 0.05, \*\**P* < 0.01).

Alpha-smooth muscle actin (αSMA) staining revealed a partial loss of signal within the medial layer in aged aortas, indicating vascular smooth muscle cell (VSMC) phenotypic switching (Fig. 4d). Correspondingly, staining for ionized calcium-binding adapter molecule 1 (IBA1) revealed an abundance of macrophages, particularly in plaque regions (Fig. 4e)^19^. Increased oxygen demand within atherosclerotic plaques often leads to angiogenesis, a source of intra-plaque hemorrhage^20^. To assess plaque progression, Prussian blue staining of iron, a component of hemosiderin in hemorrhagic processes, was performed (Fig. 4f). Quantification revealed almost no hemorrhage in young aortas, but significantly higher deposits in aged ones (Fig. 4j).

Moreover, von Kossa staining confirmed calcification in all aged aortas, to the exclusion of young ones (Fig. 4g). Hematoxylin and eosin (H&E) staining demonstrated a significant increase in medial intimal wall thickness (IMT) in aged aortas (Fig. 4h, Supplementary. Fig. 3b)^21^. To contextualize our findings for translational comparisons, plaques were classified according to the grading scheme by Renu Virmani^22^. Unlike in young animals, grading in aged animals exhibited intimal thickening and xanthomas (Fig. 4k, Supplementary Fig. 3c–f).

Interestingly, H&E staining revealed adventitial accumulations of basophilic cells in two aged animals, raising the question whether arterial tertiary lymphoid organs (ATLO) are present in the common marmoset (Fig. 4l). ATLO are accumulations of various lymphoid and other immune cells at the adventitial side of atherosclerotic plaques, which play significant roles in immune response^23,24^. Indeed, immunostaining of cluster of differentiation 3 (CD3) revealed an abundance of T-lymphocytes within the accumulations (Fig. 4l).

The aforementioned findings prompted further examination of possible underlying molecular changes in hepatic expression of genes involved in cholesterol metabolism. Indeed, expression of *ABCA1* and *ABCG5* encoding ATP-binding cassette transporters responsible for cholesterol efflux, was found decreased in aged livers (Fig. 4m–p)^25^. Interestingly, expression of *CD36*, important for lipoprotein uptake, was also significantly decreased (Fig. 4q)^26^. Expression of cholesterol 7-alpha-hydroxylase (*CYP7A*), encoding the rate-limiting enzyme in bile acid synthesis, was decreased, while no differences were found in 7-dehydrocholesterol reductase (*DHCR7*) or low-density lipoprotein receptor (*LDLR*) genes (Fig. 4r–t).

### Molecular signature of the aged heart by snRNA-sequencing

To elucidate the molecular underpinnings of the functional and morphological findings in aged hearts, we performed cardiac snRNA-sequencing (Fig. 5). Cluster annotation was similar between the age groups (Supplementary Fig. 4a). Interestingly, epicardial mesothelial cells displayed the most differentially expressed genes (DEGs) of all cell types in both age groups (Supplementary Fig. 5a). In aged hearts, these were overrepresented in pathways of fatty acid uptake, fibroblast growth factor receptor 4 (FGFR4), platelet-derived growth factor receptor and its downstream effector, PI3K, whereas the cardioprotective and pro-regenerative *IL4R* was downregulated (Fig. 5c)^27,28^. Aged cardiomyocytes upregulated stress marker genes for natriuretic peptides, *NPPA* and *NPPB*, the hypertrophy-associated Wnt ligand, *WNT16*, as well as the mitogen activated protein kinase *MAPK10*, a key driver of cardiac remodeling and fibrosis (Supplementary Fig. 5d)^29–31^. Aged arterial vEC upregulated genes involved in antigen processing and presentation, such as *B2M*, encoding the β chain of major-histocompatibility class I molecules (MHC-I), and *CD74*, recently shown to mediate EC injury response and macrophage foaming^32,33^. The class B scavenger receptor gene, *CD36*, central to atherogenesis by driving lipid-uptake mediated endothelial dysfunction, was upregulated in aged arterial vEC (Supplementary Fig. 5f)^34,35^. Interestingly, *KLF3*, an early response gene to EC inflammatory insult was also upregulated^36^. Young-adult capillary vEC upregulated genes involved in proliferation, migration, angiogenesis, barrier regulation and homeostatic flow response, including *SLIT3*, *HIF1A*, *CARMIL1* and *SMAD6*, respectively (Supplementary Fig. 5h)^37–39^. Their aged counterparts, on the other hand, upregulated a number of genes associated with EC injury response, apoptosis, endothelial dysfunction and coronary pathology, including *CD74*, *TIMP3* and *TOX2*, respectively^33,40–42^. Gene set enrichment analysis (GSEA) of capillary vEC highlighted terms of active barrier regulation in young-adult, whereas aged capillaries DEGs enriched for terms of ER stress and endothelial remodeling (e.g. EIF2AK4)^43,44^. Overrepresented terms in aged cardiac fibroblasts were those of extracellular matrix (ECM) deposition and ossification, whereas their younger counterparts overrepresented NO-cGMP signaling (Fig. 5c). While aged pericytes’ DEGs overrepresented connective tissue development, those of young-adult pericytes were overrepresented in heparan sulfate glycosaminoglycans (HS GAG) biosynthesis as well as beta-adrenergic receptor activity, reflecting greater vascular support, maturation and relaxation (Fig. 5c)^45,46^. In aged macrophages, among the most upregulated genes was plexin A4 (*PLXNA4*), an amplifier of Toll-like receptor inflammatory signaling^47^. Moreover, aged macrophages upregulated pro-inflammatory genes for phosphodiesterase 7B (*PDE7B*), and neuron navigator 2 (*NAV2*), implicated in tumor necrosis alpha (TNFα) secretion and downstream signaling, respectively^48,49^.

**Figure 5:**
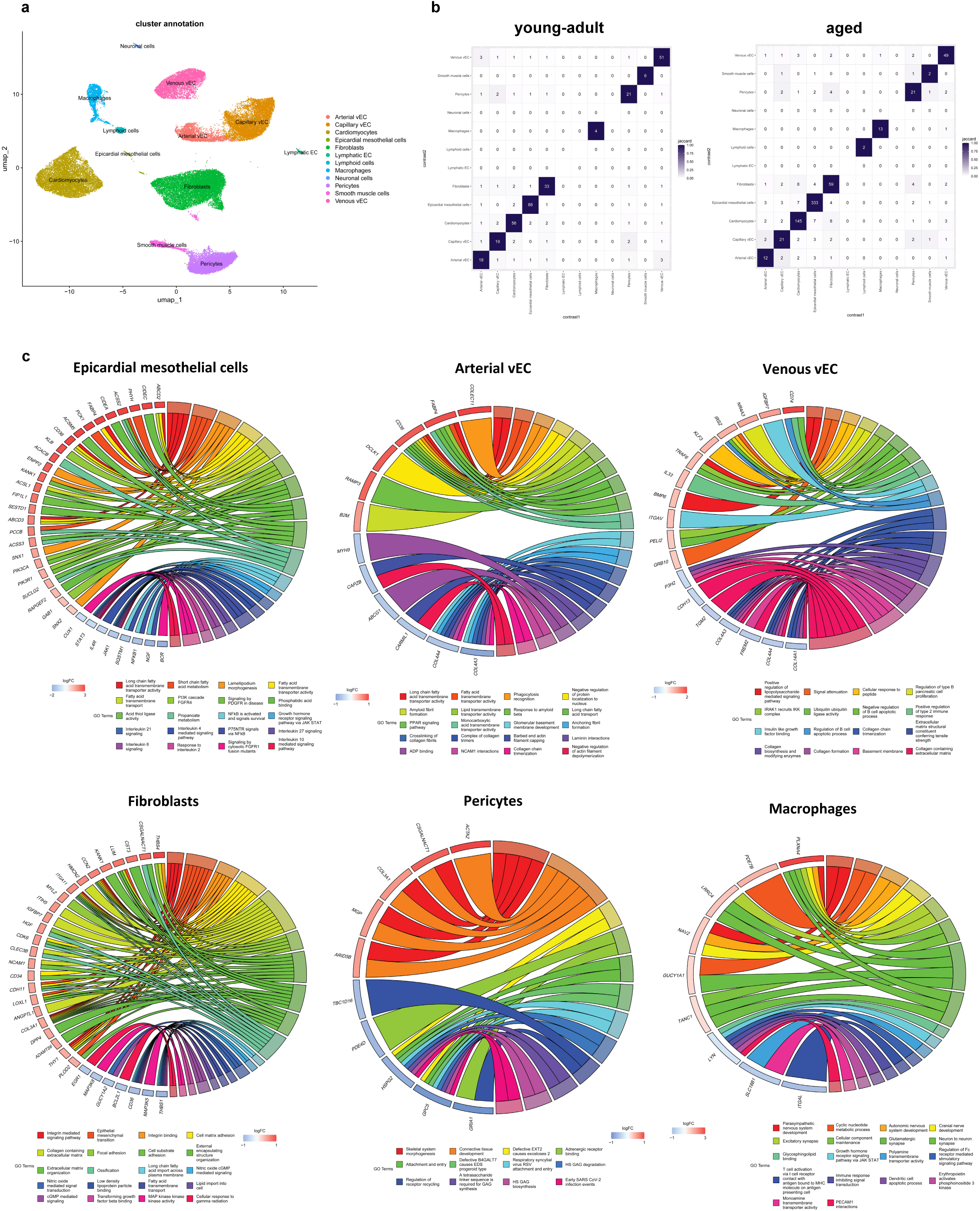
Single nuclei RNA sequencing of young-adult and aged marmoset hearts. **a**, Uniform manifold approximation and projection (UMAP) embedding of 55,392 nuclei delineates spatial clustering of 12 distinct cardiac cell types from young-adult (n = 7) and aged animals (n = 7). **b**, Jaccard plots shows the count of differentially upregulated genes (adjusted *P* value < 0.05, log2FC > 0.5) across cardiac clusters (jaccard score). **c**, Chord plots depict over representation term analysis for Gene Ontology (biological process, molecular function and cellular component), Reactome, Hallmarks and KEGG of the differentially expressed genes (adjusted *P* value < 0.05) in the aged (log2FC > 0.5; red) and young-adult (log2FC < –0.5; blue) clusters: epicardial mesothelial cells, arterial and venous vascular endothelial cells (vEC), fibroblasts, pericytes and macrophages.

To investigate intercellular communications between different cardiac clusters upon aging, we conducted CellChat analysis. There were slightly greater collective number of inferred interactions as well as interaction strength in young-adult compared to aged hearts (Supplementary Fig. 6a). Interestingly, in young-adult hearts, cardiomyocytes were the most receptive cell population, receiving the most and strongest signals from all cardiac populations, including autocrine signaling (Fig. 6a). In aged hearts, however, the major source of inferred interactions were fibroblasts, transmitting 28 signals to capillary vEC alone. Capillary vEC featured as the main receivers of signals in the aged heart including of their own (Fig. 6a, Supplementary Fig. 6b).

**Figure 6:**
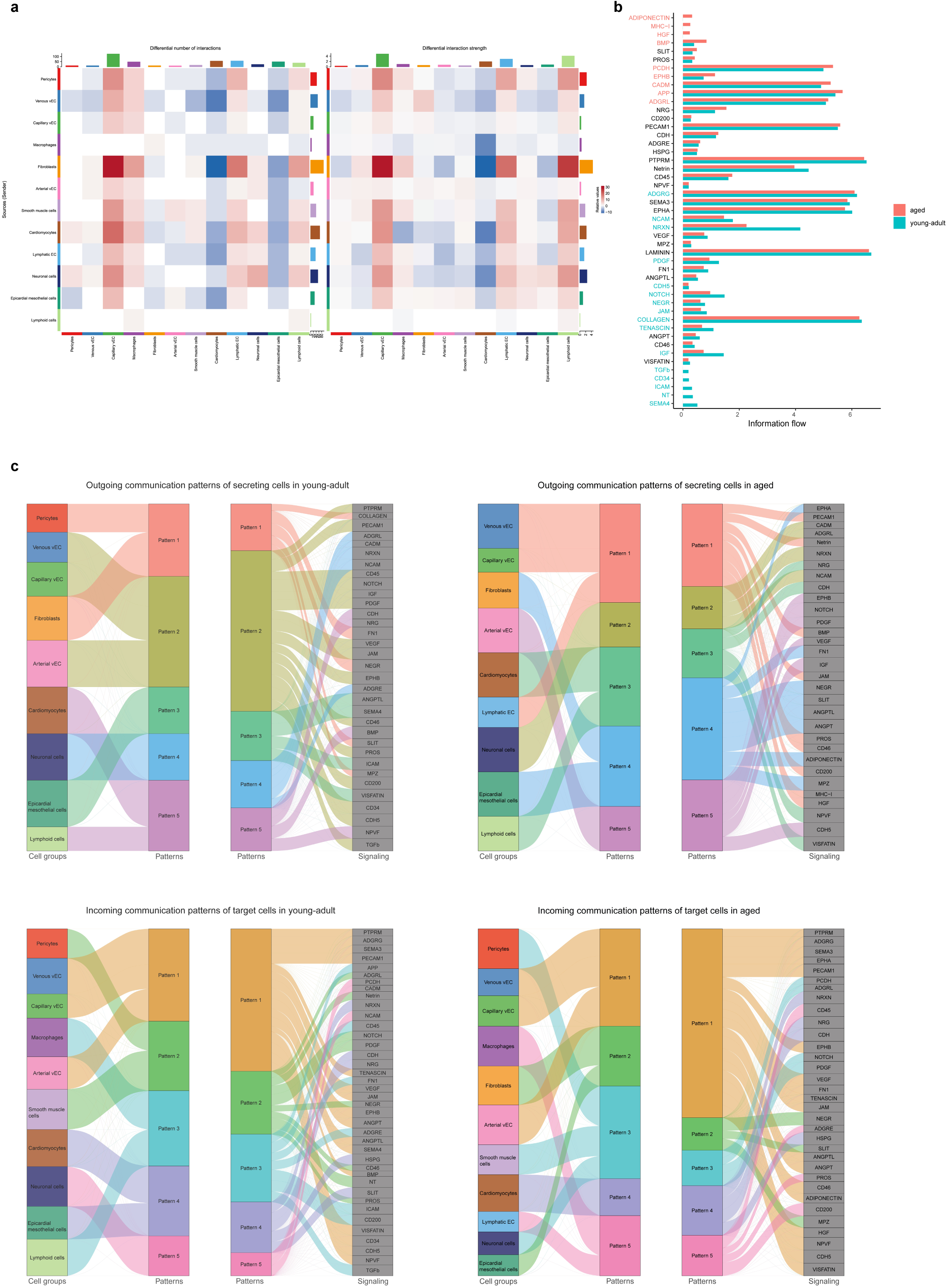
CellChat cardiac cell interaction analysis based on expression patterns revealed by single nuclei RNA sequencing. **a**, Heat map of cell interaction patterns of differentially expressed genes in young-adult (blue) or aged (red) cardiac clusters showing the number of interactions and interaction strength. **b**, Predicted information flow of different pathways. **c**, Sankey plots of outgoing and incoming cell communication patterns of different cardiac cell clusters in young-adult and aged animals.

Molecular information flow analysis highlighted a number of unique as well as shared pathways with differential enrichment between the two age groups. Pathways unique to young-adult hearts include Semaphorin 4 (SEMA4) and Neurotrophin (NT), whereas aged-exclusive ones featured Adiponectin, MHC-I and hepatocyte growth factor (HGF) (Fig. 6b).

Young-adult vEC transmitted TGFβ signals to macrophages, while receiving SEMA4 signals from the epicardium. Outgoing signals in aged hearts, however, displayed a segregation of endothelial cells, where venous, capillary and lymphatic EC converged on a pattern characterized by junctional adhesion molecule (JAM), BMP, MHC-I and HGF, whereas arterial vEC had their own pattern, i.e. EphB. Aged fibroblasts and epicardial mesothelial cells shared the same outward pattern featuring adiponectin, while aged cardiomyocytes transmitted visfatin. The incoming communication landscape further highlighted vEC groups as the most receivers of signals in the aged heart from fibroblasts and epicardium as well as inter-cellular.

### Age-associated pathogenic protein cargo of cardiac EVs contrast a cardioprotective signature in the young

EVs play a crucial role in intercellular communication^50,51^. To investigate whether the observed cardiovascular phenotype in aged animals is reflected by their cardiac EV protein cargo, we isolated and performed proteomic analysis of cardiac EVs (Fig. 7). The results revealed differential protein cargo between the two age groups (Fig. 7c, Supplementary Fig. 7a). Of the highly enriched proteins in aged cardiac EVs were those previously reported in endothelial dysfunction, coronary artery disease, platelet activation and atherosclerosis, e.g. ERBB3, EPHA4, Latexin (LXN), plasminogen activator 2 (PLA2) and Netrin 1 (NTN1), among others^52–56^. Besides its pro-atherogenic properties, NTN1 is a senescence marker, known to recruit sympathetic fibers^57^. Importantly, the insulin-like growth factor 1 receptor, IGF1R, recently implicated in age-associated heart failure was upregulated, while the Ca^+2^-binding glycoprotein, Sarcalumenin (SRL), known to decline in aging hearts was severely depleted in aged cardiac EVs^58,59^. Proteins reported in cardiac injury, ferroptosis and mitochondrial damage were also enriched in aged cardiac EVs, including the ectonucleotidase ENPP1, the 15-lipoxygenase (ALOX15) and Vitronectin (VTN), respectively^60–62^. Importantly, the NADase, CD38, was upregulated, while the rate-limiting enzyme for NAD+ salvage pathway, NAMPT, was downregulated in aged cardiac EVs compared to young-adults’, reflecting an age-associated cardiac NAD+ depletion^63,64^. The senescence-marker, CLIC3 chloride channel, as well as the pro-inflammatory and cardiac adverse remodeling prognostic marker, AXL tyrosine kinase, were both upregulated in aged cardiac EVs^65,66^. Notably, glucose transporters, SLC2A1 (GLUT1) and SLC2A4 (GLUT4) were both significantly upregulated therein (Supplementary Fig. 7a).

**Figure 7:**
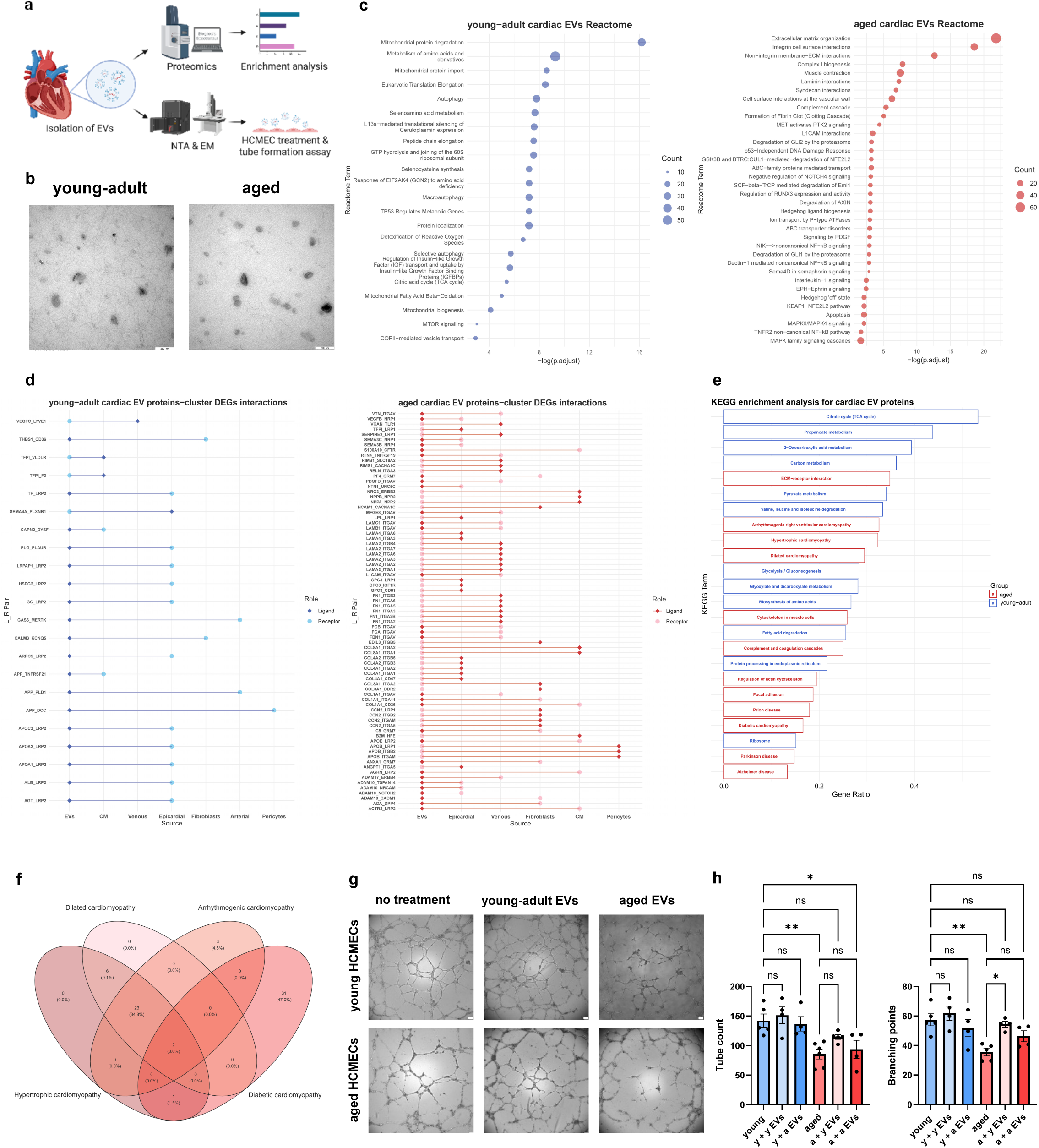
Proteomic and functional characterization of cardiac extracellular vesicles (EVs). **a**, Workflow (figure created in BioRender). **b**, Exemplary electron microscopy images of cardiac EVs derived from young-adult and aged animals. Scale bar = 250 nm. **c**, Reactome pathway enrichment analysis displaying unique terms for differentially expressed EV proteins in young-adult and aged animals (n = 5). **d**, Expected ligand-receptor (L_R) interaction in differentially expressed cardiac EV proteins and DEGs of different cardiac clusters (sn-RNA sequencing) in young-adult and aged animals in reference to CellTalkDB. **e**, KEGG pathway enrichment analysis of differentially expressed cardiac EV proteins from young-adult and aged animals. **f**, Venn diagram of cardiomyopathy-associated proteins from KEGG enrichment analysis of aged-upregulated cardiac EV proteins. **g**,**h**, Tube formation assay in young and aged HCMECs treated with cardiac EVs from young-adult or aged animals showing bright field images of tube networks (scale bar = 100 μm) (**g**) and ImageJ quantification of tube count and branching points (n = 4–5) (**h**). Bar graphs represent means ± SEM. Statistical significance was assessed by one-way ANOVA and Šídák’s multiple comparisons test (\**P* < 0.05, *\*\*P* < 0.01, ns: not significant).

Reactome pathway enrichment analysis of young-adult cardiac EV protein cargo highlighted a number of cardioprotective pathways of mitochondrial vigilance, active redox system, autophagy and proteostasis (Fig. 7c, Supplementary Fig. 7a). The aged Reactome, on the other hand, featured a prominent enrichment for ECM deposition and interactions, reflecting a pro-fibrotic trajectory (Fig. 7c, Supplementary Fig. 7a). Herein, terms with pro-inflammatory, pro-thrombotic, vascular pro-adhesive and atherogenic properties were strongly represented. Notably, enriched terms for degradation of the hedgehog downstream effectors GLI1 and GLI2, along with “Hedgehog ‘off’ state” might imply a failed cardiac regenerative capacity in aged hearts^67,68^. DNA damage responses, either p53-dependent or independent, are further hallmarks of senescence enriched by cardiac aged EV proteins^69^. Finally, activation of MAPK corroborates an augmented pro-hypertrophic signaling in aged hearts (Fig. 7c).

To infer possible molecular interactions between cardiac EV proteins and different cardiac cell populations, we integrated cardiac EV proteomic and snRNA-seq analyses in reference to the ligand-receptor interaction database (CellTalkDB)^70^. In young-adult hearts, ligands were predominantly enriched in EVs. Of these, the apolipoproteins, APOA1 and APOA2, known for their protective cardiometabolic associations, partnered with the epicardially expressed lipoprotein-associated receptor LRP2^71^. Importantly, the young-adult cardiac arterial EC-expressed TAM receptor MER tyrosine kinase (MERTK) finds its canonical ligand, GAS6, in the EVs, an interaction known to induce EC migration, angiogenesis and downstream phosphorylation of endothelial nitric oxide synthase (eNOS)^72^. Potential interactions in aged hearts, however, were dominated by ECM proteins, which find receptors among EV-enriched integrins. Interestingly, the heart failure correlated marker of activated fibroblasts dipeptidyl-peptidase-4 (DPP4), expressed in aged cardiac fibroblasts finds its ligand adenosine-deaminase (ADA) in the EVs^73,74^. Importantly, the aged epicardially upregulated neuropilin receptor, NRP1, potentially pairs with the EV-enriched class 3 semaphorins, SEMA3B and SEMA3C; an embryonically crucial interaction known to be reactivated in response to cardiac injury^75,76^. Interestingly, aged cardiomyocytes upregulate beta-2-microglobin (B2M), which is reported to spike upon ischemic damage and activates cardiac fibroblasts^77^. It finds its interacting partner, hereditary hemochromatosis protein (HFE) enriched in the EVs; a complex crucial for cellular iron uptake, with evidence linking cardiac iron overload and fibrosis^78,79^.

To gain an overview of the functional patterns of EV proteins, Proteomaps were created (Supplementary Fig. 7b)^80,81^. The young-adult cardiac EV cargo was highly represented in metabolic and proteostatic activity, whereas the aged featured maladaptive terms, e.g. PI3K-Akt signaling. Moreover, KEGG pathway analysis of aged cardiac EV protein cargo featured significant enrichments for cardiomyopathies, as well as inflammation, remodeling, atherosclerosis and thrombosis (Fig. 7e,h, Supplementary Fig. 7c,d).

As our results demonstrated an age-driven impairment in cardiac microvascular supply, we sought to evaluate the functional relevance and translational potential of cardiac EVs in this regard. To this end, primary HCMECs from young or old donors were treated with cardiac EVs from young-adult or aged marmosets, followed by tube formation assays. At baseline, older HCMECs displayed significantly diminished angiogenesis (Supplementary Fig. 4c–i). Interestingly, when treated with young-adult marmoset cardiac EVs, tube formation was significantly enhanced, implicating a pro-angiogenic potential of young EVs (Fig. 7j–l; Supplementary Fig. 4k).

### Aged aortas transcriptomics reveal early signals of vascular pathology

To further elucidate the molecular patterns in diseased vessels, bulk RNA-sequencing was performed in atherosclerotic (aged) and lesion-free (young-adult) abdominal aortas (Figure 8a, Supplementary Fig. 5a–e).

**Figure 8:**
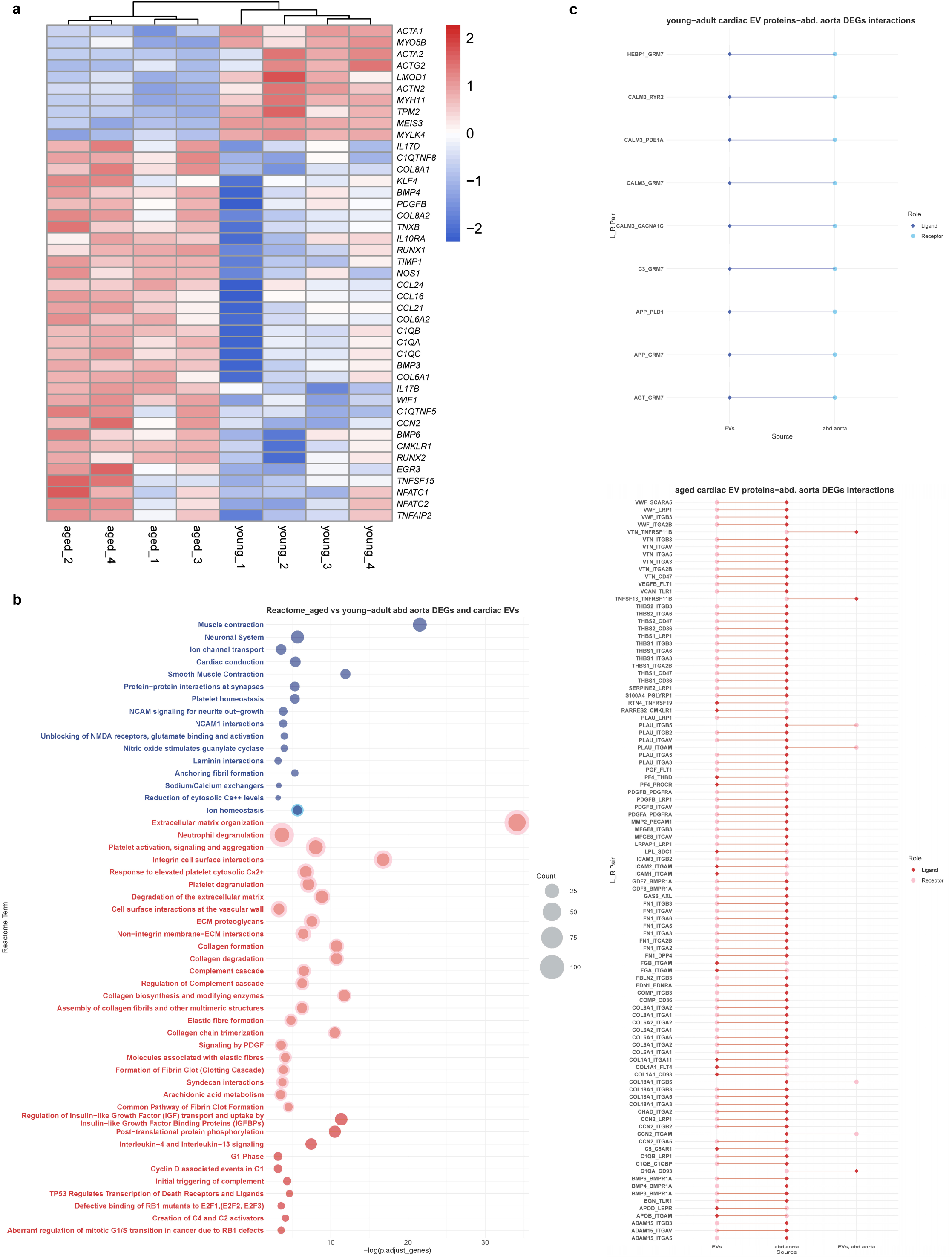
Bulk RNA sequencing of abdominal aortas of common marmosets and predicted interaction with cardiac EVs. **a**, Heat map showing a selection of differentially expressed genes from young-adult and aged animals (n = 4). **b**, Reactome pathway enrichment analysis showing unique terms for DEGs in abdominal aortas and those commonly shared by cardiac EV proteins in each age group. **c**, Expected ligand-receptor (L_R) interaction in DEGs of abdominal aorta from young-adult and aged animals and differentially expressed cardiac EV proteins from corresponding age groups in reference to CellTalkDB.

Aged abdominal aortas upregulated expression of transcription factors characteristic of VSMC migration, phenotypic switching, inflammation and calcification (Fig. 8a). These include nuclear factors of activated T-cells (*NFATC1*, *NFATC2*), Runt-related transcription factors (*RUNX1*, *RUNX2*) and cellular communication network factor 2 (*CCN2*)^82–85^. Herein, upregulation of osteoblast-like differentiation marker genes, such as bone morphogenic proteins (*BMP4*, *BMP6*) and tissue inhibitor of metalloproteinase (TIMP1) coincided with downregulation of the VSMC contractile markers, α– and γ-actins (*ACTA1*, *ACTA2*, *ACTG2*), β-tropomyosin (*TPM2*), leiomodin 1 (*LMOD1*), myosin heavy chain (*MYH11*) and myosin light chain kinase (*MYLK4*)^82,86^. Interestingly, T-cell recruitment and activation markers were also upregulated in aged aortas, including chemokine ligands (*CCL16*, *CCL21* and *CCL24*) and platelet derived growth subunit B (*PDGFB*)^82,87–89^. Notably, macrophage complement family genes (*C1QA*, *C1QB*, *C1QC*), as well as nitric oxide synthase (*NOS1*) were also upregulated^90,91^. Finally, the Wnt inhibitor factor (*WIF1*) was among the top upregulated genes in aged aortas.

Reactome pathway enrichment analysis highlighted a number of unique terms in lesion-free vs. atherosclerotic DEGs. Integration with the cardiac EV proteomics dataset displayed some shared terms, highlighting interaction continuities. For example, young-adult abdominal aorta’s DEGs were enriched for terms of vascular homeostasis, e.g. smooth muscle contraction, nitric oxide synthesis and ion homeostasis. The latter was shared by the young-adult cardiac EV protein Reactome (Fig. 8b). The aged, on the other hand, featured a number of shared and unique terms with overwhelmingly pathological connotations, including ECM deposition, proliferation, calcification and thrombosis (Fig. 8b).

In an attempt to dissect the aforementioned term analysis, we inferred ligand-receptor interaction between and within DEGs in abdominal aortas and cardiac EV proteins. Aged interactions featured key molecular pairings involved in vascular pathological progression. For example, the cardiac EV-loaded VEGF receptor, FLT1, met its ligand, VEGFB, in the aorta, likely promoting intra-plaque angiogenesis. Moreover, the TNF receptor superfamily 11B (TNFRSF11B, or osteoprotegrin), involved in EC dysfunction and vascular inflammation, finds potential partners, including VTN and the TNF superfamily 13, TNFSF13, a marker of atherosclerosis and platelet activation. Thrombospondins 1 and 2, both markers of aortic aneurysm, expressed in aged aortas, act as ligands to EV-packaged receptors, e.g. CD36, CD47. Finally, a number of ECM ligand-receptor pairs are inferred between the two aged compartments, reflecting the role of cardiac EVs in augmenting fibrosis-driven vascular remodeling.

## Discussion

The term ‘radical life extension’ refers to the unprecedented increase in human longevity since the twentieth century^92^. Not only has humanity made revolutionary medical triumphs over morbid ailments, but also achieved an astonishing level of public health awareness of ‘healthy aging’. However, even in the absence of comorbidities, natural aging entails gradual organ system decrepitude, albeit with subclinical manifestations. Here, we sought to investigate the impact of aging on the cardiovascular system independent of comorbidities, utilizing the common marmoset as a NHP model.

While conventional non-invasive cardiovascular functional diagnostic measures showed little to no deviations in healthy aged animals, only upon PV loop measurements that a deterioration in cardiac function was detected. Notably, we previously demonstrated the superior diagnostic utility of PV loop compared to MRI^93^. Moreover, histological examination revealed a conspicuous cardiovascular phenotype in aged animals, characterized by interstitial and perivascular fibrosis, microvascular rarefaction, as well as cardiomyocytes size and structural aberrations. These findings highlight general shortcomings of conventional methods in reflecting early age-driven cardiovascular changes. Molecular investigations confirmed the histopathological findings, pointing towards impaired cholesterol metabolism, endothelial dysfunction, fibroblast activation, mild chronic inflammation and cardiomyocyte stress and pro-hypertrophic signaling. Importantly, cardiac intercellular communication revealed a loss of cardiomyocyte centrality in the aged heart, whereas young cardiomyocytes remained a hub of incoming signals from all cardiac cell types.

Discrepant expression patterns of the cholesterol scavenger receptor, CD36, between the aged liver and cardiac vEC indicate an age-associated lipid dislocation. Moreover, the microvascular phenotype reported in our study confirms the vulnerability of the microcirculation at the onset of cardiovascular pathology, particularly in the context of aging^94^. Age-associated cardiac EC dysfunction was also accompanied by pericyte aberrant gene programs with strong implications for loss of vascular tone, phenotype switching and increased ECM deposition, which might be responsible for the observed perivascular fibrosis. Indeed, Tamiato et al. identified a pro-fibrotic phenotype of senescent pericytes contributing to cardiac fibrosis and reduced LV-EF^95^. Interestingly, young-adult cardiac pericytes’ gene program reflected an intimate connection to vEC through glycocalyx recycling and tight barrier regulation.

Aged cardiac macrophages were primed to inflammation, upregulating genes known to aggravate, rather than initiate, an inflammatory response. Their young-adult counterparts, however, engaged in TGFβ signaling, known to induce M2 polarization and suppress inflammation^96^.

Aged cardiac fibroblasts upregulated nearly twice as many genes as in young-adults and dominated outward signals of all cardiac cell types, both in number and strength, reflecting their activated state. The most receptive of these signals in the aged heart were capillary vEC, potentially explaining the observed perivascular fibrosis and highlighting the importance of fibroblast-microvascular EC communication in age-driven cardiac fibrosis. Importantly, cellular crosstalk revealed an involvement of aged cardiac fibroblasts in adiponectin signaling. While adiponectin signaling is thought to be protective in certain instances of severe cardiac insults, it comes with a cost of increased collagen remodeling and degradation of elastin, potentially leading to impaired cardiac relaxation^97^. In contrast, young-adult cardiac fibroblasts activated NO-cGMP signaling, which has been experimentally shown to communicate to cardiomyocyte β-receptor cAMP signaling, conferring a homeostatic regulation of cardiac lusitropy^98,99^. These findings sketch the early molecular steps on the path to age-driven cardiac fibrosis and ventricular stiffness, revealing a transition to diastolic dysfunction corroborated by our PV loop measurements. While no differences were observed between the age groups in stroke work or cardiac output, systolic function impairment typically ensues later on^100,101^.

Interestingly, epicardial cells dominated the DEG count by snRNA-sequencing analysis, highlighting an important molecular contrast in this cardiac compartment upon aging worthy of further investigations. Herein, upregulated fatty-acid transporters in the aged epicardium might exacerbate the observed cardiovascular pathology^102^. In this study, we leveraged cardiac intercellular crosstalk upon aging by proteomic investigation of cardiac EVs. Unsurprisingly, young cardiac EVs were loaded with cardioprotective and anti-oxidant proteins associated with mitochondrial fitness and proteostasis. Aged cardiac EVs, on the other hand, enriched for senescence-associated, NAD-depleting, pro-fibrotic, pro-inflammatory and pro-hypertrophic proteins. Interestingly, diabetic cardiomyopathy was most prominently represented by aged cardiac EV proteins. Therein, upregulated glucose transporters reflect an early metabolic transition to heart failure, where increased glucose utilization compensate for deteriorated mitochondrial function^103^. In the absence of diet or genetic confounders, these results underscore aging alone as a driver of cardio-metabolic syndrome. Moreover, our study highlights for the first time a number of proteins that have not been known to be secreted in cardiac EVs, providing a potential route of their cardiac activity. For example, the CYP450 POR, has previously been shown to confer protection against cardiomyocyte hypertrophy and remodeling if expressed in cardiac ECs, albeit with unexplained mechanism^104^. Here, we show that POR is enriched in cardiac EVs from young animals (Supplementary Fig. 7a). While the cardiac cell population responsible for its EV-shedding is unidentified in our study, future research should consider the role of cardiac EV-enriched proteins in maintaining cardiac homeostasis.

Moreover, we provide the first experimental evidence of proangiogenic properties of young cardiac EVs, evoking their translational potential to reverse capillary rarefaction in old hearts. Indeed, our proteomic analysis of young-adult cardiac EVs revealed a number of proteins with pro-angiogenic and vascular network stabilizing properties, e.g. the guanylate exchange factor, Trio (Supplementary Fig. 7a)^105^.

In congruence with histological findings, analysis of bulk RNA-seq data from aged animals’ abdominal aortas indicate an age-associated molecular signature of vascular inflammation, VSMC transdifferentiation and loss of contractile apparatus. Prominent examples here include genes for NFATC, RUNX1 and RUNX2 transcription factors^106^. These three have been reported as downstream effectors of CD8+ T-cell-mediated plaque progression, VSMC osteogenic differentiation and loss of contractility in mice models of atherosclerosis^82^. Importantly, recent single-cell transcriptomic studies examining cross-species divergence in atherosclerosis highlighted the T-cell-dominated adaptive immune component in human lesions compared to mouse^107^. In line with these studies, our data show upregulated signature genes of T-cell attractants in aged marmoset aortas, thereby leveraging the human translatability of our model. Intriguingly, *WIF1*, found significantly upregulated in aged aortas aligns with an earlier report by Zhang et al. describing this Wnt inhibitor as a primate-specific marker of arterial aging confined to aortic EC, with yet unidentified roles^108^.

Furthermore, *CCN2* identified in our bulk RNA-seq of lesioned aortas is a pro-fibrotic factor upregulated in hypoxic EC by visfatin^109–111^. CellChat analysis predicted visfatin to be secreted by aged cardiomyocytes, suggesting an interplay between the heart and vessels, and encouraging future investigations of the visfatin-CCN2 axis.

In the context of atherosclerosis, EV cargo can drive plaque progression^112^. Herein, our integrative analysis of cardiac EV proteomic and aortic RNA-sequencing allude to EVs’ potential in linking molecular, structural, and functional changes in both the myocardium and vessels and highlight a number of ligand-receptor pairs as potential therapeutic targets subject to future investigations.

Our NHP model displayed an aging phenotype with great resemblance to that in humans^113–118^. Moreover, the common marmoset exhibits a similar lipoprotein profile to humans and has been suggested as an excellent model for lipoprotein metabolism and diet-induced atherosclerosis^119,120^. Indeed, previous reports found different plaque stages in aged marmosets^121^. To the contrast, conventional rodent models do not naturally develop adverse lipid profiles or atherosclerosis during aging, thereby limiting their human translatability^122–125^. Our findings in the aged common marmoset strongly suggest that age alone is a sufficient risk factor for atherosclerosis development and progression^8,126^.

As human cardiovascular aging is often accompanied by comorbidities, it is crucial for precision medicine to distinguish disease patterns emanating from these comorbidities from those solely due to aging^127^. The common marmoset, free from confounding factors, characterized for the first time in our study presents itself as a powerful translational model for human aging research.

## Materials & Methods

### Animals

The animals were raised and kept at the German Primate Center, Göttingen, Germany. All aspects of animal husbandry and experimental procedures adhered to the European Union Directive 2010/63/EU and the German Animal Welfare Act. The animal experiment protocol received approval from the institutional animal welfare committee and was subsequently sanctioned by the Lower Saxony State Office for Consumer Protection and Food Safety (LAVES, number of approval: 33.19-42502-04-20/3458). All experimental procedures complied with the relevant regulations of German law. All animals in this study were kept and treated in a species-appropriate manner and received veterinary care whenever necessary.

Marmosets were housed indoors in steel cages with a partner animal or in a family group. The cages were equipped with custom-made sleeping boxes, hammocks, natural branches, wood chips as bedding, and weekly changing equipment such as swings or boxes for enrichment purposes. The room temperature was kept constant at 26 ± 1 °C, with a relative humidity of 60-80 % and 10 air changes per hour. Circadian rhythms were maintained with alternating 12-hour periods of light and darkness. The diet consisted of a portion of porridge with curd or milk mash and additional vitamins, minerals and proteins in the morning. At noon, the animals received a mixture of fruit, vegetables and pellets (ssniff® Mar, ssniff Spezialdiäten GmbH, Soest, Germany). As a contribution to enrichment, the feeding schedule was slightly changed daily, e.g. by adding mealworms, gum Arabic, nuts, seeds or eggs. Water was offered *ad libitum* in drinking bottles.

### Experimental design

Ten young and ten old male common marmosets were chosen randomly by the husbandry administration of the German Primate Center. Animals aged two to four years were considered “young-adult”, while animals over eleven years were considered “aged” in this study, meeting published aging criteria in this animal species^128^. The median life span of common marmosets in captivity is approximately 10 years. The sample size calculation using G*Power software, based on the primary age-related statistical characteristics from a previous study^6^, indicated that a group size of 10 animals is required for the left ventricular end-systolic volume (LV ESV) in one-sided tests. For the other relevant parameters (left ventricular end-diastolic volume, right ventricular end-diastolic and end-systolic volumes), smaller group sizes were determined.

Before incorporating the animals into the study, a clinical and hematological assessment was conducted to ensure only clinically healthy animals were included. Heart function was acquired using MRI, followed by invasive intra-cardiac pressure-volume loop measurements. An interval of minimum four weeks was maintained between the two functional cardiac measurements to mitigate any potential impact of the MRI procedure on the subsequent hemodynamic pressure-volume measurement. This precaution additionally ensured a sufficient recovery of the animals after the MRI anesthesia.

Body weight was acquired prior to each procedure performed on the animals. Subsequently after the PV loop measurements, animals were euthanized and a broad spectrum of organ samples was collected for further analyses.

For histological analyses, samples from the German Primate Center’s collection of further healthy animals’ tissue sections were included to increase the sample size provided that they met the same criteria for age group allocation.

Whenever direct intervention with the animal was necessary, such as anesthesia for functional cardiac measurements, blood sampling, or sample collection during necropsy, investigators were aware of the animal’s identity and allocation to the respective age group. This ensured correct animal identification and preparedness for potential age-related anesthetic incidents. During data analysis, samples were handled and analyzed in a blinded manner whenever feasible.

### Hematological assessment

Blood was collected via the femoral artery using a 26G cannula. The volume of blood collected ranged from 1.5 to 2 ml per animal, adhering to the recommendations of the Society for Laboratory Animal Science (GV-SOLAS; “Recommendation for blood collection from laboratory animals, especially small laboratory animals,” 2017). A complete blood count was performed using the Advia 2120 Hematology System (Siemens Healthcare Diagnostics Inc., Tarrytown, NY, USA). Serum profile analysis was conducted by the Institute for Clinical Chemistry of University Medical Center of Göttingen (UMG).

### Cardiac MRI

Cardiovascular MRI was conducted using a 9.4 T MR system (BioSpec 94/30, Bruker BioSpin, Germany) equipped with a 330 mT/m gradient system (BGA-20S, Bruker BioSpin, Germany).

All imaging studies were performed using a custom-made ellipsoid single-loop receive coil with a diameter of 38 mm × 35 mm (O-HLE-94, Rapid Biomedical, Germany), which was positioned underneath the chest for optimal heart coverage. Functional CMRI was performed using a navigator based IntraGate-FLASH sequence during free breathing in short-axis view with following parameters: 40 navigator points, repetition time (TR) = 10.5 ms, echo time (TE) = 1.6 ms, flip angle = 35°, Field of View (FOV) = 51.2 × 51.2 mm2, Matrix size = 256 × 256, echo position = 33.33%, spatial resolution = 0.2 × 0.2 × 1.2 mm3, 2 × 2 in-plane interpolation, slice gap = 0.3 mm, 12 slices, and cardiac phases = 16. Left and right ventricle functions were assessed using short-axis images via the freely available software package Segment (Version 2.0 R6435, Medviso, Lund, Sweden). The end-diastolic volume (EDV) and end-systolic volume (ESV) were calculated using Simpson’s method of disks. Left-ventricular stroke volume and ejection fraction (EF) were computed as per the equations:

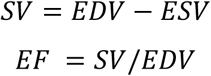

First-pass perfusion MR imaging was performed after an intravenous dose of 0.125 mmol per kilogram of body weight of Gd-DTPA (Gadovist; Bayer Schering Pharma AG, Berlin, Germany) using a non-ECG gated, golden angle, radial FLASH sequence (TR = 6 ms, TE = 1.5 ms, flip angle = 5°, FOV = 32 × 32 mm2, echo position = 50%, spatial resolution = 0.5 × 0.5 × 2.5 mm³, number of spokes = 137, number of repetition = 150) including a saturation recovery module. Myocardial perfusion ratio (MPR) was assessed using the ratio of the signal-time curve slope in myocardium and blood during the first pass of the contrast agent.

T1 mapping of the left ventricle before and after administration of the contrast agent was performed in a mid-ventricular short-axis slice using inversion recovery non-ECG gated golden angle radial FLASH without breath holding (TR = 4.5 ms, TE = 2.1 ms, flip angle = 5°, FOV = 32 × 32 mm2, echo position = 50%, spatial resolution = 0.5 × 0.5 × 2 mm³, number of spokes = 5000, and number of repetition = 3).

The extracellular volume (ECV) was calculated using the following equation: *ECV* = (1 − *hct*) × (Δ*R*1*myo*/Δ*R*1*blood*) where R1 = 1/T1. A hematocrit (hct) is an average of all hct values taken after each CMR study.

### Hemodynamic pressure-volume measurement

Intra-cardiac hemodynamic measurements and data analysis was performed as previously described^93^. Briefly, animals were put under deep general anesthesia. Vital parameters were acquired continuously. Since the abdominal approach was chosen for the intervention, the abdominal cavity and the diaphragm were opened to expose the heart. Blood and muscle resistivity were measured and entered to the ADV500 system. A loose suture was put around the vena cava inferior for vena cava occlusions to gain load independent hemodynamic measures. The PV-loop catheter was inserted into the heart and the measurement was started at baseline heart frequency. Vena cava occlusions were conducted by carefully lifting the thread around the vein. After baseline and occlusion data were collected, the heart frequency was increased by application of dobutamine (0.9-14.4 ml/kg/h, careful infusion until heart rate was increased by 30-50%). After all data were collected, blood was drawn for final analyses, animals were euthanized under deep anesthesia with an overdose of potassium chloride (>75 mg/kg BW, Kaliumchlorid 7.45 %, B.Braun, Germany) and underwent immediate necropsy.

### Sampling

Organs were collected during the necropsy. After excision and weighing of the heart, the ventricles and atria were separated from each other. Each heart sample was sectioned following the same scheme. The tissue sections were either directly processed, fixated in 4% formaldehyde, or brought to liquid nitrogen and stored at – 80°C, depending on the location (left/right atrium, left/right ventricle) and the respective analysis.

Thoracic aortas were excised and stored in 4 % paraformaldehyde. The anterior half of the abdominal aorta was placed in 10 % formaldehyde, the caudal half was frozen to –80° C. The aorta sampling scheme is illustrated in the Supplementary Figure 3a. Liver samples were brought to liquid nitrogen directly after excision.

### Multiphoton microscopy

Waviness analysis of myosin content of the LV was performed using label-free Second Harmonic Generation (SHG) measurements as previously described in Khan et al., 2021^15^. Briefly, SHG imaging was conducted on 50 µm vibratome tissue sections using an upright TriM Scope II 2-Photon microscope (Miltenyi Biotec, Bielefeld, Germany) equipped with a tunable femtosecond laser with two independently tunable output channels (Cronus 2P, Light Conversion, Vilnius, Lithuania). The first tunable channel (680 – 960 nm) was used for SHG excitation. The incident laser power under the objective was 19 mW at 870 nm. Images were acquired using a Zeiss W Plan-Apochromat 20x (NA 1.0) water immersion objective. A PMT (Hamamatsu) in the transmission position collected the forwardscattered emitted light gathered by a 1.4 NA condenser lens under the stage. SHG signals were collected using the filter 434±20 nm (BrightLine HD filters, Semrock, AHF Analysentechnik Tübingen, Germany). 2D mosaics of entire sections were acquired with individual image sizes of 336 × 336 µm with 1024 × 1024 pixels, a pixel dwell time of 2.19 µs with 2-fold averaging and 10% overlap within each tile of the mosaic. Magnified images of regions of interest (ROI) were acquired with a 112 × 112 μm field of view and 512 × 512 pixels, ensuring sufficient resolution for analysis. The ROIs were selected within the myocardial layer of the left ventricular and interventricular septum (IVS) walls. The magnified SHG images were used for measuring the waviness of the myofibrils to assess their hypercontractile state (Fig. 2o,p). Individual fibers were manually traced using ImageJ. The traced fibers were then segmented with custom Python script using the OpenCV library. For each fiber, the arc length and straight length were measured, and the waviness was calculated as the ratio of straight length to arc length. A higher ratio indicated straighter fibers, while a lower ratio indicated more waviness.

A detailed assessment of vascular supply in the hearts was performed using label-free SHG, third-harmonic generation (THG) and 3-photon autofluorescence measurements by 3-photon excitation microscopy. Briefly, 100 µm vibratome tissue sections were imaged using a TriM Scope Matrix multiphoton microscope (Miltenyi Biotec) equipped with a wavelength-tunable laser source (1300 – 1700 nm, CRONUS 3P, Light Conversion, Vilnius, Lithuania). The incident laser power under the objective was ≈ 5mW at 1300 nm. Images were acquired using a 25x Olympus (NA 1.05) water immersion objective. The backscattered emitted light was split by two 495 nm and 590 nm long pass 2” dichroic mirrors (Semrock) and detected at photomultiplier tube (PMT) detectors (Hamamatsu). The forward scattered emitted light gathered by a 1.4 NA condenser lens under the stage was split by a 495 nm long pass 1” dichroic mirror (Semrock) and detected at the PMT in the transmission position. SHG and THG signals were collected in forward scatter using the filter setting 647 nm long pass and 432±36 nm (AHF Analysentechnik AG) while 3-photon autofluorescence was collected in backscatter. 3D mosaics of entire heart transverse sections were acquired with individual image sizes of 294 × 294 µm with 1024 × 1024 pixels and 10% overlap within each tile of the mosaic. THG signals of elastic fibers inside of blood vessels were used for quantification of the vascular supply in the heart samples.

### Histology heart

For Masson Trichrome staining, 3 µm tissue sections of the left ventricle were fixed in 4% formaldehyde and paraffin-embedded. Then, descending alcohol series were performed in a staining machine (DiaPath Giotto, HistoServe, Celle, Germany). Sections were incubated in Bouin’s solution overnight at room temperature. Bouin’s solution was removed by carefully rinsing the slides in aqua dest before the following steps were carried out: 2 min Weigert’s hematoxylin – 10 min running tap water – 5 min azophloxine – 30 seconds 1 % acetic acid – 2 min phosphoromolybdic acid orange – 30 seconds 1 % acetic acid – 5 min light green – 5 min 1 % acetic acid. After this, microscope slides were carefully dried sparing the tissue. Slides were then brought to 96 % ethanol for 2 min. This was followed by two 2 min washing steps in 100 % ethanol and three 2 min washing steps in xylene. After this step, slides were ready for mounting with Eukitt (Sigma-Aldrich/ Merck, Darmstadt, Germany, #03989500).

Images of the slides were taken using a laser microscope (Leica DMi8, Leica Microsystems CMS GmbH, Wetzlar, Germany). For each localization within the heart (left and right atrium, left and right ventricle), five randomly selected images per animal at a 40× magnification were included in the evaluation of interstitial fibrosis.

### Immunofluorescence heart

Wheat germ agglutinin (WGA) staining was performed to determine cardiomyocyte cross-sectional area. Detection of infiltration with inflammatory cells was enabled by immunofluorescence staining with an antibody for cluster of differentiation 45 (CD45), a pan-leucocyte marker.

For immunofluorescence staining, 8 µm cryosections of the left ventricle were used. Briefly, sections were fixed in acetone and washed in PBS. Blocking was performed to avoid unspecific bindings, the antibody against CD45 (Purified anti-marmoset CD45, Clone 6C9, BioLegend, San Diego, CA, U.S.) was applied and samples were incubated overnight. After another washing step, samples were incubated with the secondary antibody (AlexaFluor™, 594 goat anti-mouse IgG (H+L), invitrogen, Waltham, MA, U.S.). For the WGA-staining a conjugated antibody was used, making this step void (WGA, Alexa Fluor® 488 conjugate, invitrogen). Again, slides were washed and then mounted with a mounting medium containing DAPI for staining of nuclei (VECTASHIELD® Antifade Mounting Medium with DAPI, Vector Laboratories, Newark, CA, U.S.).

Images were taken one day after the staining process with a laser microscope in 40 x magnification (Leica DMi8, Leica Microsystems CMS GmbH, Wetzlar, Germany).

Analysis was carried out using ImageJ. For WGA staining, cells were marked via free hand selection and the area of each cell was calculated by the program. CD45 signals were marked via color threshold and the area was calculated by the program. At least five randomly selected images per animal were analyzed.

### Single-nuclei isolation and RNA-sequencing

The nuclei isolation protocol was carried out as described before^129^. After the nuclei extraction from cardiac tissues, the nuclei were stored in nuclei storage buffer, thereafter washed and resuspended in 1ml of 2 % BSA/PBS for sorting via DAPI-staining. Finally, 10,000 DAPI-positive single nuclei were collected in 2 % BSA/PBS solution (Becton-Dickinson FACSAria III Fusion) for sequencing.

Library preparation for single nuclei mRNA-Seq analysis was performed according to the Chromium NextGEM Single Cell 3ʹ Reagent Kits v3.1 User Guide (Manual Part Number CG000204 Rev B; 10x Genomics) with 14 cycles for cDNA amplification. 10,000 cells were loaded to the 10x controller in order to reach a target number of 5,000 cells per sample. Fragment length distribution of generated libraries was monitored using ‘Bioanalyzer High Sensitivity DNA Assay’ (5067-4626; Agilent Technologies). Quantification of libraries was performed by use of the ‘Qubit® dsDNA HS Assay Kit’ (Q32854; ThermoFisher Scientific).

The generated mRNA expression libraries were pooled accordingly, denatured with NaOH, and were finally diluted to 1.8 pM according to the ‘Denature and Dilute Libraries Guide’ (Document # 15048776 v02; Illumina). 1.3 ml of denatured pool was sequenced on an Illumina NextSeq550 sequencer using one high output flowcell for 75 cycles and 400 million clusters (#20024906; Illumina). Sequencing was performed according to the following settings: 28bp as sequence read 1; 56bp as sequence read 2; 8bp as index read 1; no index read 2.

The 10x Genomics CellRanger analysis pipeline set (v7.0.0) was used with default parameters except for the setting of expected cells (5,000 cells) and did not allow any barcode mismatches. Default CellRanger quality filtering were performed and the CellRanger count pipeline was used to align red data to the reference genome provided by 10X Genomics (Callithrix_jacchus.mCalJac1.pat.X release-106 (Ensemble)). Single-nucleus RNA-sequencing (snRNA-seq) data were processed using the Cell Ranger pipeline with default parameters to generate gene–barcode count matrices.

Downstream analyses were performed in R using Seurat (v5.0.0). Data were merged into a single Seurat object prior to quality control. Low-quality nuclei were excluded based on gene complexity thresholds (nFeature_RNA > 500, nFeature_RNA < 4500). Data normalization and principal component analysis (PCA) were conducted using Seurat default settings. Dimensionality reduction was performed using Uniform Manifold Approximation and Projection (UMAP), and nearest-neighbor graphs were constructed using the first 20 principal components.

Clustering was performed at a resolution of 0.2. Cell type annotation was done using the FindAllMarkers function together with canonical marker gene expression. Differential gene expression analyses were performed within each cell type using the FindMarkers function with default parameters. Genes were considered differentially expressed with cut-off abs (avg_log2FC) > 0.5 and adjusted p value (p_val_adj) < 0.05. Pairwise overlaps among differentially expressed gene sets were computed using a custom R script.

Functional enrichment analyses were performed using the clusterProfiler package (v4.14.6) using Hallmark, KEGG, Reactome, and Gene Ontology (GO) pathways. Enrichment results were visualized as chord diagrams using the circlize package (v0.4.16). Gene set enrichment analysis was performed using fgsea (v1.32.4), and genes ranked by avg_logFC values and evaluated against the same pathway collections.

Cell–cell communication analyses were done using CellChat (v2.1.2) with default parameters applying the default human ligand-receptor database. CellChat objects were created for each condition, and communication probabilities and pathways were computed^130^.

### Extracellular vesicle isolation and purification

During necropsy, roughly 300 mg of heart tissue from the left ventricle were given to perfusion buffer: 135 mM NaCl, 4 mM KCl, 1 mM MgCl_2_, 10 mM HEPES, 0.33 mM NaH_2_PO_4_, 10 mM glucose, 10 mM 2,3-butanedione monoxime (Sigma, St Louis, MO), 5 mM taurine (Sigma, St Louis, MO) and pH adjusted to 7.2. 150-200 mg of the tissue were weighed and given to a petri dish. The heart tissue was minced for 5 min with a scalpel, then moved to a falcon with digestion buffer: 0.5 mg/ml Liberase TL (Roche Diagnostics, Mannheim, Germany), 20 µg/ml DNase (Qiagen, Hilden, Germany), 10 mM HEPES, DMEM, followed by incubation at 37°C for 15 min in a water bath. The tube was inverted every minute to help dissociation. With a 5 ml pipet, the digestion was resuspended and filterated with a 40 µm cell stainer, using the plunger of a 1 ml syringe. The filter was washed with 500 µl of DMEM. The suspension was centrifuged 330g for 6 min at 4°C. The supernatant was collected and centrifuged at 2000g for 20 min at 4°C. The supernatant was then collected and either frozen at –20°C or directly used for the purification.

For purification, the supernatant was transferred to a 6.5 ml tube for ultracentrifugation 16000g for 20 min at 4°C to remove detritus. The supernatant was again brought to a new tube and ultracentrifuged 118000 g for 90 min at 4°C. The remaining supernatant was sucked and the pellet was resuspended in 500 µl cold PBS. 1 ml of 60% Iodixanol was added and mixed gently. 2 ml of 30% Iodixanol were layered carefully on top. Finally, a layer of 1200 µl 10% Iodixanol was carefully added. The border between the 30% and 10% Iodixanol layers was labeled. The tube was then ultracentrifuged 186000g for 2.5 hours at 4°C. The interphase between the 30% and the 10% Iodixanol layer was collected and transferred to a fresh tube with 3 ml cold PBS and centrifuged 118000 g for 1 hour at 4°C for washing. Afterwards, the supernatant was discarded and the remaining pellet was either resuspended in 30 µl 4x Laemmli + DTT for Proteomics or resuspended in 50 – 100 µl PBS (depending on pellet size) for nanoparticle tracking analysis and subsequent cell culture^131^.

For freezing the EVs without a significant loss of function, EVs were stored in a PBS-HEPES-Albumin-Trehalose Buffer at –80°C^132^.

### Nanoparticle tracking analysis & electron microscopy

To determine the concentration of the cardiac extracellular vesicles, a nanoparticle tracking analyzer was used (Nanosight NS300, Malvern Panalytical Ltd, Malvern, UK). For the measurement, the EV solution was diluted 1:100 in PBS. Five measurements á 1 min were performed per sample and the mean content was calculated (Nanosight NTA 3.4, Malvern Panalytical Ltd, Malvern, UK). The solution was saved for electron microscopy which was performed according to Théry et al., 2006 to ensure quality of isolation^133^.

### Proteomics of the extracellular vesicles

Protein samples were purified by running them a short distance into a 4 – 12% NuPAGE Novex Bis-Tris Minigel (Invitrogen), visualized by Coomassie staining, and prepared for bottom-up proteome analysis by in-gel digestion with trypsin^134^.

Protein digests were analyzed on a nanoflow chromatography system (Dionex nanoRSLC, Thermo Fisher Scientific) hyphenated to a hybrid timed ion mobility-quadrupole-time of flight mass spectrometer (timsTOF Pro 2, Bruker). In brief, 400 ng equivalents of peptides were dissolved in loading buffer (2% acetonitrile, 0.1% trifluoroacetic acid in water) and spiked with a synthetic peptide standard used for retention time alignment (iRT Standard 1:100 (v/v), Schlieren, Switzerland).

Digest samples were enriched on a reversed-phase C18 trapping column (0.3 cm × 300 µm, Thermo Fisher Scientific) and separated on a reversed-phase C18 column with an integrated CaptiveSpray Emitter (Aurora 25 cm × 75 µm, IonOpticks) using a 100 min linear gradient of 5 – 35 % acetonitrile/0.1% formic acid (v:v) at 250 nl/min, at a column temperature of 50⁰C.

DIA analysis was performed in diaPASEF mode using a custom 20×2 variable window isolation scheme with windows from m/z 400 to 1,200 to include the 2+/3+/4+ population in the m/z–ion mobility plane. The collision energy was ramped linearly as a function of the mobility from 59 eV at 1/K0 =1.5 Vs cm−2 to 20 eV at 1/K0=0.7 Vs cm−2. Three technical replicates per biological replicate were acquired.

Protein identification was achieved by directDIA processing with the Pulsar algorithm in Spectronaut Software 16.0 (Biognosys) using default settings against the UniProtKB *Callithrix jacchus* reference proteome v1.2023 (48.340 entries) augmented with a set of 53 known common laboratory contaminants. For quantitation, up to the 6 most abundant fragment ion traces per peptide, and up to the 10 most abundant peptides per protein were integrated and summed up to provide protein area values. Mass and retention time calibration as well as the corresponding extraction tolerances were dynamically determined. Both identification and quantification results were trimmed to a False Discovery Rate of 1% using a forward-and-reverse decoy database strategy. Multiple testing correction was performed by Storey-Tibshirani method to determine Q-values. Analysis of differential protein candidates (average log2 ratio > 0.5 or < –0.5 and cutoff threshold of Q-value < 0.05) was performed in RStudio. Scatter plot, pathway enrichment analysis in the Kyoto Encyclopedia of Genes and Genomes (KEGG), Reactome pathway enrichment analysis and their corresponding plot representations were performed using the R packages clusterProfiler, org.Hs.eg.db, enrichplot, ReactomePA, ggplot2 and ggvenn.

### Ligand_Receptor interaction inference

Possible ligand-receptor interactions between the EV proteomics and RNA sequencing datasets were inferred using CellTalkDB as a reference. CellTalkDB provides a manually curated and literature-supported ligand-receptor interaction (LR pairs) database^70^. Differentially enriched cardiac EV proteins in young-adult or aged animals were integrated with DEG lists of cardiac clusters of snRNA sequencing data or with bulk RNA sequencing data of abdominal aortas (see below) from the two age groups. CellTalkDB human_lr_pair.rds was used as a reference to find ligand-receptor pair matches in the datasets and the analysis was performed in RStudio.

### Cell culture

Primary human cardiac microvascular endothelial cells (HCMEC; C-12286; PromoCell, Heidelberg, Germany) obtained from cardiac ventricles of three Caucasian males aging at 15 (young), 50 or 51 years old were cultured in PromoCell microvascular media (MV; PromoCell C-22020) supplemented with corresponding supplement mixes (PromoCell C-39225 or C-39226, respectively) and 0.1% penicillin/streptomycin (PS). HCMEC were maintained in a humidified incubator at 37°C Celsius and 5% CO_2_ and used for experiments between passages 4 to 6. The cells were treated with EVs derived from either young or aged marmosets’ hearts for 48h at a concentration of 1×10^8^ per mL medium. Angiogenesis by tube formation assay was performed in μ-Slide Angiogenesis (#81506, Ibidi GmbH, Gräfelfing, Germany). The cells were seeded on Matrigel matrix basement membrane (Corning 354234) at 12,000 cells/well. At 7 hours, bright-field images were taken using Leica DMi8 microscope (Leica Microsystems CMS GmbH, Wetzlar, Germany). Tube formation was analyzed by Image J Angiogenesis Analyzer software plugin and total tube count per image was considered as angiogenesis parameter.

### Oil Red O staining aorta

En Face Oil Red O (ORO) staining of the thoracic aorta was performed based on the protocol of Andrés-Manzano *et al.*^17^ with some modifications. Briefly, after necropsy thoracic parts of the aortas were fixed in 4 % paraformaldehyde for one day (aorta sampling scheme: Suppl. Fig. S3A). 0.2 % Oil Red O solution was prepared by adding 0,07 g Oil Red O powder (Santa Cruz Biotechnology Inc., Dallas, Texas, USA) to 25 ml 100 % methanol and 10 ml NaOH and mixing thoroughly. Right before usage, the solution should be filtered twice through a 0.45 µm filter. Prior to the staining, aortas were cleaned and adhesive tissue was removed. Samples were washed in 78 % methanol twice for 5 min, followed by 50 min incubation in the Oil Red O solution at room temperature on a tilted roller. Afterwards, aortas were transferred to a clean tube and repeatedly washed in 78 % methanol twice for 5 min. For imaging, the vessel was opened longitudinally following the great curvature by the help of micro scissors and micro forceps. With small needles, the vessel was fixed on a black ground and images were taken using a stereomicroscope (STEMI 508 and Axiocam 208 color, both ZEISS, Oberkochen, Germany) in combination with the Zen blue software (ZEN blue Version 3.4.91.00000, Carl Zeiss Microscopy GmbH). Image analysis was performed using ImageJ (ImageJ 1.53k, Wayne Rasband and contributors, National Institutes of Health, USA).

### Histology aorta

After the ORO staining, 3 µm sections of formalin-fixed paraffin-embedded tissue were prepared. Hematoxylin eosin (HE) staining was performed using a staining machine (DiaPath Giotto, HistoServe, Celle, Germany), showing no degradation of tissue quality caused by the previous ORO staining.

HE stained slides were used to measure the average wall thickness of each animal’s thoracic aorta. This was done by selecting the whole tissue cross section and calculating the average thickness in Leica Application Suite X program (Leica Microsystems GmbH, Wetzlar, Germany).

Masson Trichrome staining was performed as described above to determine collagen content.

Elastica van Gieson staining to detect organization of elastic fibers in the aortic tissue started with a descending alcohol series, followed by 20 min incubation in Elastica solution. Slides were then washed with running tap water and aqua dest for 5 min, respectively, and afterwards put to Weigerts hematoxylin for 2 min. After a washing step of 10 min in running tap water, slides were brought to van-Gieson-solution for 5 min. In the end, slides were dehydrated and mounted with Eukitt mounting medium (Sigma-Aldrich/ Merck, Darmstadt, Germany). Collagen content and elastic fiber content were set into relation to calculate the elastin/collagen ratio per animal.

von Kossa staining was done starting with descending alcohol series in the stainer (DiaPath Giotto, HistoServe, Celle, Germany), followed by 45 – 60 min incubation in silver nitrate solution. Slides were reduced 2 min in sodium carbonate formaldehyde solution and washed with running tap water, then incubated for 5 min in 5 % sodium thiosulphate solution. After another step of washing with tap water and aqua dest, slides were brought to nuclear fast red for 5 min and were again washed. The staining was completed after ascending alcohol series in the stainer and the final mounting of the slides with Eukitt mounting medium. The percentage content of calcified material per vessel slice was calculated.

Prussian Blue staining started with descending alcohol series in the stainer (DiaPath Giotto, HistoServe, Celle, Germany), then 5 min in 10 % potassium ferrocyanide. This was followed by a 30 min incubation in hydrochloric acid-potassium ferrocyanide and a washing step in distilled water. Afterwards, slides were brought to nuclear fast red for 5 min and again washed. The ascending alcohol series in the stainer was followed by mounting with Eukitt. Plaque hemorrhage was quantified in each slide.

Atherosclerotic lesion classification was performed according to Virmani et al. 2000^22^. Briefly, lesions are morphologically classified in 7 categories consistent with the American Heart Association categorical scheme for “intermediate” or “advanced” plaques (i.e. type IV, V, and VI lesions). These are intimal xanthoma, intimal thickening, pathological intimal thickening, fibrous cap atheroma, thin fibrous cap atheroma, calcified nodule, and fibrocalcific plaque.

For the analysis of histologic staining, Leica Application Suite X program (Leica Mycrosystems GmbH, Wetzlar, Germany) was used.

### Immunohistochemistry aorta

Immunohistochemistry (IHC) with antibodies for ionized calcium-binding adapter molecule 1 (Iba1 antibody, Cat No. GTX101495, GeneTex, Irvine, CA, USA) and alpha smooth muscle actin (Clone 1A4, DakoCytomation Denmark A/S, Glotrup, Denmark) as well as CD3 (Dako M0851, Agilent, Santa Clara, CA, USA) was performed in an immunohistochemistry staining machine (Discovery XT, Ventana, Roche Diagnostics GmbH, Mannheim, Germany) to detect macrophages, smooth muscle cells, and T-lymphocytes, respectively, in the plaque tissue.

### RNA isolation

For RNA-Isolation, 20 mg of liver or aortic tissue, respectively, was disaggregated (gentleMACS™ Octo Dissociator and corresponding M-Tubes, Miltenyi Biotec, Bergisch Gladbach, Germany) and RNA was isolated by using NucleoSpin® miRNA kit (Macherey & Nagel, Dueren, Germany). RNA quantification and quality control was performed by spectrophotometry (NanoDrop™ – Thermo Scientific); guidelines regarding wave length absorbance ratio of 260/280 and 260/230 were followed, so that the ratios between the wave lengths 230:260:280 of 1:2:1 were accepted.

### Quantitative PCR (qPCR)

Liver: 1000 ng RNA were used for cDNA synthesis by Omniscript® Reverse Transcription Kit (Qiagen, Hilden, Germany). Quantitative PCR was carried out using TaqMan™ Fast Advanced Master Mix (Applied Biosystems, Waltham, MA, USA) and the following TaqMan® Gene Expression Assays (primers): *ABCA1* (Cj01059115_m1), *ABCG1* (Cj06129309_s1), *ABCG5* (Cj01043228_m1), *ABCG8* (Cj06064440_m1), *CD36* (Cj05946203_m1), *CYP7A1* (Cj06124698_m1), *DHCR7* (Cj06066729_m1), *HMGCR* (Cj05940336_m1) and *LDLR* (Cj05949905_m1).

Succinate dehydrogenase complex subunit A served as an endogenous control (*SDHA*, Cj04399031_m1) (PMID: 23451040). All primers were purchased from ThermoFisher Scientific, Waltham, MA, U.S.

Real-Time PCR (RT-PCR) was run according to the recommended thermal cycling profiles of the StepOnePlus™ and software v2.3 (Thermo Scientific) to calculate threshold cycles (C_T_). Relative quantification of gene expression was calculated according to the 2*^−ΔΔCT^* formula, using C_T_ values for *SDHA* as housekeeping control.

### Bulk RNA-sequencing (abdominal aorta)

RNA isolated from 8 marmoset aortas underwent bulk sequencing as follows: RNA quality was assessed by measuring the RNA integrity number (RIN) using a Fragment Analyzer HS Total RNA Kit. Library preparation for RNA-Seq was performed on the STAR Hamilton NGS automation platform using the Illumina Stranded mRNA Prep, Ligation, and the Illumina RNA UD Indexes Set A, Ligation, starting from 300 ng of total RNA. The size range of the final cDNA libraries was determined with the SS NGS Fragment 1-6000 bp Kit on the Fragment Analyzer, with an average size of 340 bp. Quantification of the cDNA libraries was performed using the QuantiFluor™ dsDNA System (Promega). Libraries were amplified and sequenced on an S4 flow cell NovaSeq 6000, 100 cycles, generating 25 million reads per sample (Illumina). Sequencing images were processed using BaseCaller Illumina software to generate BCL files, which were demultiplexed into fastq files with bcl2fastq v2.20.0.422.

Raw read & Quality check: Sequence images were transformed with Illumina software BaseCaller to BCL files, which was demultiplexed to FastQ files with bcl2fastq v2.20. Sequencing quality was asserted using FastQC (Babraham Bioinformatics).

Mapping & Normalization: Sequences were aligned to the reference genome *Callithrix jacchus* (mCalJac1, https://www.ensembl.org/Callithrix_jacchus/Info/Index) using the RNA-Seq alignment tool (version 2.7.10) allowing for 2 mismatches within 50 bases^135^. Subsequently, read counting was performed using featureCounts^136^. Read counts were analyzed in R/Bioconductor environment (version 4.3.1, www.bioconductor.org) using the DESeq2 package version 1.40.2^137^. Candidate genes were filtered using an absolute log2 fold-change > 0.5 (aged) and < –0.5 (young-adult) and FDR-corrected adjusted P value < 0.05. A heat map for selected DEGs was created in RStudio using pheatmap R package. Reactome pathway enrichment analysis and representation (cutoff adjusted P value < 0.05) was created in R studio using the R packages ReactomePA and ggplot2.

### Statistics

Statistics were performed using GraphPad Prism (GraphPad Software, Boston, MA, U.S.). Comparison of two groups assuming Gaussian distribution was conducted in Student’s t-Tests. P values < 0.05 were considered significant. Multiple comparisons assuming Gaussian distribution were compared using one-way analysis of variance (ANOVA) with Šídák’s multiple comparisons test.

### Data availability

Raw and pre-processed cardiac sn-RNA sequencing are deposited in NCBI’s Gene Expression Omnibus (GEO)^138^ and are accessible through GEO Series accession number GSE319494 (https://www.ncbi.nlm.nih.gov/geo/query/acc.cgi?acc=GSE319494, the following secure token has been created to allow review of record GSE319494 while it remains in private status: cdwnkqewfdgnlkr).

Raw and normalized sequencing data for bulk RNA sequencing of abdominal aortas are deposited in GEO under accession number GSE317015 (https://www.ncbi.nlm.nih.gov/geo/query/acc.cgi?acc=GSE317015, the following secure token has been created to allow review of record GSE317015 while it remains in private status: mnofscwunlchvgx).

The mass spectrometry proteomics data have been deposited to the ProteomeXchange Consortium via the PRIDE^139^ partner repository and are available with the dataset identifier PXD073658 (access token: H1nY4znLDhhf).

All data will be publicly released upon publication.

## Author contributions

R.H., M.M. and G.G. designed the study. L.K. and M.M. performed *in vivo* experiments including physiological characterization, PV loop measurements and analysis. A.M., T.R.M., M.R., J.K., L.K., M.M. and S.B. performed MRI measurements and analysis. L.K. and M.M. performed and analyzed histological findings. L.K. performed RNA isolation and qPCR. L.K., G.G. and C.L. isolated and analyzed EVs *ex vivo*. M.Sa., G.G. and L.K. analyzed *in silico* proteomic data. M.Sa. interpreted EV proteomic data. M.Sa. performed and anlyzed EVs *in vitro* experiments. J.L.Y., D.L., M.Sa., F.B., C.B., M.K., L.Z. and T.T. performed snRNA-seq including bioinformatics. M.Sa. interpreted snRNA-seq results. M.Sa. performed snRNA-seq and L_R interaction analysis and interpretation. F.R.G., A.Kh., A.Ku. and F.A. performed and analyzed label-free histology. M.Si., G.S. performed bulk RNA-sequencing of the abdominal aorta and M.Sa. analyzed interpreted the data. W.M. and B.S. performed electron-microscopy and quality control of the EVs. S.S. advised and interpreted atherosclerosis and liver data. R.B. provided study concept advice. L.K., M. Sa., G.G., M.M. and R.H. wrote the manuscript. All authors commented on and edited the manuscript.

## References

1 Ciumărnean, L. et al. Cardiovascular Risk Factors and Physical Activity for the Prevention of Cardiovascular Diseases in the Elderly. Int J Environ Res Public Health 19 (2021). 10.3390/ijerph19010207

2 North, B. J. & Sinclair, D. A. The intersection between aging and cardiovascular disease. Circulation research 110, 1097–1108 (2012). 10.1161/circresaha.111.246876

3 Triposkiadis, F., Xanthopoulos, A. & Butler, J. Cardiovascular Aging and Heart Failure: JACC Review Topic of the Week. Journal of the American College of Cardiology 74, 804–813 (2019). 10.1016/j.jacc.2019.06.053

4 Senos, R. et al. Gross morphometry of the heart of the Common marmoset. Folia Morphol (Warsz) 73, 37–41 (2014). 10.5603/fm.2013.0064

5 Falcão, B. M. R. et al. Heart anatomy and topography of the common marmoset (Callithrix jacchus Linnaeus, 1758). J Med Primatol 49, 153–157 (2020). 10.1111/jmp.12463

6 Moussavi, A. et al. Cardiac MRI in common marmosets revealing age-dependency of cardiac function. Sci Rep 10, 10221 (2020). 10.1038/s41598-020-67157-5

7 Ross, C. N., Davis, K., Dobek, G. & Tardif, S. D. Aging Phenotypes of Common Marmosets (Callithrix jacchus). J Aging Res 2012, 567143 (2012). 10.1155/2012/567143

8 Fischer, K. E. & Austad, S. N. The development of small primate models for aging research. Ilar j 52, 78–88 (2011). 10.1093/ilar.52.1.78

9 Tardif, S. D., Mansfield, K. G., Ratnam, R., Ross, C. N. & Ziegler, T. E. The marmoset as a model of aging and age-related diseases. Ilar j 52, 54–65 (2011). 10.1093/ilar.52.1.54

10 Drummer, C. et al. Generation and Breeding of EGFP-Transgenic Marmoset Monkeys: Cell Chimerism and Implications for Disease Modeling. Cells 10 (2021). 10.3390/cells10030505

11 Sasaki, E. et al. Generation of transgenic non-human primates with germline transmission. Nature 459, 523–527 (2009). 10.1038/nature08090

12 Kuehnel, F. et al. Parameters of haematology, clinical chemistry and lipid metabolism in the common marmoset and alterations under stress conditions. J Med Primatol 41, 241–250 (2012). 10.1111/j.1600-0684.2012.00550.x

13 Schipke, J. et al. Assessment of cardiac fibrosis: a morphometric method comparison for collagen quantification. Journal of applied physiology (Bethesda, Md.: 1985) 122, 1019–1030 (2017). 10.1152/japplphysiol.00987.2016

14 Aboonabi, A. & McCauley, M. D. Myofilament dysfunction in diastolic heart failure. Heart failure reviews 29, 79–93 (2024). 10.1007/s10741-023-10352-z

15 Khan, A. et al. Label-free imaging of age-related cardiac structural changes in non-human primates using multiphoton nonlinear microscopy. Biomed Opt Express 12, 7009–7023 (2021). 10.1364/boe.432102

16 Masi, S. et al. Assessment and pathophysiology of microvascular disease: recent progress and clinical implications. European heart journal 42, 2590–2604 (2021). 10.1093/eurheartj/ehaa857

17 Andrés-Manzano, M. J., Andrés, V. & Dorado, B. Oil Red O and Hematoxylin and Eosin Staining for Quantification of Atherosclerosis Burden in Mouse Aorta and Aortic Root. Methods in molecular biology (Clifton, N.J.) 1339, 85–99 (2015). 10.1007/978-1-4939-2929-0_5

18 Robert, L. Aging of the vascular-wall and atherosclerosis. Experimental gerontology 34, 491–501 (1999). 10.1016/s0531-5565(99)00030-3

19 Blümm, C. et al. PAC1 deficiency reduces chondrogenesis in atherosclerotic lesions of hypercholesterolemic ApoE-deficient mice. BMC cardiovascular disorders 23, 566 (2023). 10.1186/s12872-023-03600-5

20 Parma, L., Baganha, F., Quax, P. H. A. & de Vries, M. R. Plaque angiogenesis and intraplaque hemorrhage in atherosclerosis. European journal of pharmacology 816, 107–115 (2017). 10.1016/j.ejphar.2017.04.028

21 Pezzano, A., La Carrubba, S. & Gullace, G. Carotid intima-media thickness. Journal of cardiovascular medicine (Hagerstown, Md.) 7, 555–559 (2006). 10.2459/01.JCM.0000234774.81077.a6

22 Virmani, R., Kolodgie, F. D., Burke, A. P., Farb, A. & Schwartz, S. M. Lessons from sudden coronary death: a comprehensive morphological classification scheme for atherosclerotic lesions. Arteriosclerosis, thrombosis, and vascular biology 20, 1262–1275 (2000). 10.1161/01.atv.20.5.1262

23 Gräbner, R. et al. Lymphotoxin beta receptor signaling promotes tertiary lymphoid organogenesis in the aorta adventitia of aged ApoE−/− mice. J Exp Med 206, 233–248 (2009). 10.1084/jem.20080752

24 Yin, C., Mohanta, S. K., Srikakulapu, P., Weber, C. & Habenicht, A. J. Artery Tertiary Lymphoid Organs: Powerhouses of Atherosclerosis Immunity. Frontiers in immunology 7, 387 (2016). 10.3389/fimmu.2016.00387

25 Chen, L., Zhao, Z. W., Zeng, P. H., Zhou, Y. J. & Yin, W. J. Molecular mechanisms for ABCA1-mediated cholesterol efflux. *Cell cycle (Georgetown*, Tex.) 21, 1121–1139 (2022). 10.1080/15384101.2022.2042777

26 Ramji, D. P. Growth hormone-releasing peptides, CD36, and stimulation of cholesterol efflux: cyclooxygenase-2 is the link. Cardiovascular research 83, 419–420 (2009). 10.1093/cvr/cvp195

27 Bakhshian Nik, A., Alvarez-Argote, S. & O’Meara, C. C. Interleukin 4/13 signaling in cardiac regeneration and repair. American journal of physiology. Heart and circulatory physiology 323, H833–h844 (2022). 10.1152/ajpheart.00310.2022

28 Olivey, H. E. & Svensson, E. C. Epicardial-myocardial signaling directing coronary vasculogenesis. Circulation research 106, 818–832 (2010). 10.1161/circresaha.109.209197

29 Zhao, Y. et al. An essential role for Wnt/β-catenin signaling in mediating hypertensive heart disease. Scientific reports 8, 8996 (2018). 10.1038/s41598-018-27064-2

30 Deng, Y. W. et al. Hyperglycemia promotes myocardial dysfunction via the ERS-MAPK10 signaling pathway in db/db mice. Laboratory investigation; a journal of technical methods and pathology 102, 1192–1202 (2022). 10.1038/s41374-022-00819-2

31 Liu, F. et al. Time series RNA-seq analysis identifies MAPK10 as a critical gene in diabetes mellitus-induced atrial fibrillation in mice. Journal of molecular and cellular cardiology 168, 70–82 (2022). 10.1016/j.yjmcc.2022.04.013

32 Cleary, S. J. et al. Complement activation on endothelium initiates antibody-mediated acute lung injury. The Journal of clinical investigation 130, 5909–5923 (2020). 10.1172/jci138136

33 Ke, K. et al. Increased Expression of CD74 in Atherosclerosis Associated with Inflammatory Responses of Endothelial Cells and Macrophages. Biochem Genet 62, 294–310 (2024). 10.1007/s10528-023-10421-w

34 Cho, S. CD36 as a therapeutic target for endothelial dysfunction in stroke. Current pharmaceutical design 18, 3721–3730 (2012). 10.2174/138161212802002760

35 Aguilar, V. et al. Endothelial Stiffening Induced by CD36-Mediated Lipid Uptake Leads to Endothelial Barrier Disruption and Contributes to Atherosclerotic Lesions. Arteriosclerosis, thrombosis, and vascular biology 45, e201–e216 (2025). 10.1161/atvbaha.124.322244

36 Knights, A. J. et al. Krüppel-like factor 3 (KLF3) suppresses NF-κB-driven inflammation in mice. The Journal of biological chemistry 295, 6080–6091 (2020). 10.1074/jbc.RA120.013114

37 Zhang, B. et al. Repulsive axon guidance molecule Slit3 is a novel angiogenic factor. Blood 114, 4300–4309 (2009). 10.1182/blood-2008-12-193326

38 Ullah, K. & Wu, R. Hypoxia-Inducible Factor Regulates Endothelial Metabolism in Cardiovascular Disease. Frontiers in physiology 12, 670653 (2021). 10.3389/fphys.2021.670653

39 Ruter, D. L. et al. SMAD6 transduces endothelial cell flow responses required for blood vessel homeostasis. Angiogenesis 24, 387–398 (2021). 10.1007/s10456-021-09777-7

40 Wang, Q., Song, F., Dong, J. & Qiao, L. Transient exposure to elevated glucose levels causes persistent changes in dermal microvascular endothelial cell responses to injury. Ann Transl Med 9, 758 (2021). 10.21037/atm-20-7617

41 Qi, J. H. & Anand-Apte, B. Tissue inhibitor of metalloproteinase-3 (TIMP3) promotes endothelial apoptosis via a caspase-independent mechanism. Apoptosis : an international journal on programmed cell death 20, 523–534 (2015). 10.1007/s10495-014-1076-y

42 Cheng, F. et al. Identification of Differentially Expressed Genes and Prediction of Expression Regulation Networks in Dysfunctional Endothelium. Genes 13 (2022). 10.3390/genes13091563

43 Hamada, Y., Masuda, T., Ito, S. & Ohtsuki, S. Regulatory Role of eIF2αK4 in Amino Acid Transporter Expression in Mouse Brain Capillary Endothelial Cells. Pharm Res 41, 2213–2223 (2024). 10.1007/s11095-024-03793-0

44 Zhu, M. M., Dai, J., Dai, Z., Peng, Y. & Zhao, Y. Y. GCN2 kinase activation mediates pulmonary vascular remodeling and pulmonary arterial hypertension. JCI Insight 9 (2024). 10.1172/jci.insight.177926

45 Griffin, M. E., Sorum, A. W., Miller, G. M., Goddard, W. A., 3rd & Hsieh-Wilson, L. C. Sulfated glycans engage the Ang-Tie pathway to regulate vascular development. Nat Chem Biol 17, 178–186 (2021). 10.1038/s41589-020-00657-7

46 Wu, Y. et al. Biology and function of pericytes in the vascular microcirculation. Animal Model Exp Med 6, 337–345 (2023). 10.1002/ame2.12334

47 Wen, H., Lei, Y., Eun, S. Y. & Ting, J. P. Plexin-A4-semaphorin 3A signaling is required for Toll-like receptor– and sepsis-induced cytokine storm. J Exp Med 207, 2943–2957 (2010). 10.1084/jem.20101138

48 de Medeiros, A. S. et al. Identification and characterization of a potent and biologically-active PDE4/7 inhibitor via fission yeast-based assays. Cellular signalling 40, 73–80 (2017). 10.1016/j.cellsig.2017.08.011

49 Wang, R. et al. Neuron navigator 2 is a novel mediator of rheumatoid arthritis. Cell Mol Immunol 18, 2288–2289 (2021). 10.1038/s41423-021-00696-7

50 Palazzo, C., D’Alessio, A. & Tamagnone, L. Message in a Bottle: Endothelial Cell Regulation by Extracellular Vesicles. Cancers (Basel) 14 (2022). 10.3390/cancers14081969

51 Lin, Q., He, P., Tao, J. & Peng, J. Role of Exosomes in Cardiovascular Diseases. Reviews in cardiovascular medicine 25, 222 (2024). 10.31083/j.rcm2506222

52 Bian, J.-S. et al. ErbB3 Governs Endothelial Dysfunction in Hypoxia-Induced Pulmonary Hypertension. Circulation 150, 1533–1553 (2024). doi:10.1161/CIRCULATIONAHA.123.067005

53 Khan, N. et al. Inhibiting Eph/ephrin signaling reduces vascular leak and endothelial cell dysfunction in mice with sepsis. Science translational medicine 16, eadg5768 (2024). 10.1126/scitranslmed.adg5768

54 Zhang, X. Abstract 4364291: Latexin Promotes Neointimal Hyperplasia by Enhancing SMC Proliferation and Macrophage Migration through Activation of MAPK Signaling. Circulation 152, A4364291–A4364291 (2025). doi:10.1161/circ.152.suppl_3.4364291

55 Kugiyama, K. et al. Circulating Levels of Secretory Type II Phospholipase A 2 Predict Coronary Events in Patients with Coronary Artery Disease. Circulation 100, 1280-1284 (1999). doi:10.1161/01.CIR.100.12.1280

56 Schlegel, M. et al. Silencing Myeloid Netrin-1 Induces Inflammation Resolution and Plaque Regression. Circulation research 129, 530–546 (2021). doi:10.1161/CIRCRESAHA.121.319313

57 Yu, A. Q. et al. Senescent Cell-Secreted Netrin-1 Modulates Aging-Related Disorders by Recruiting Sympathetic Fibers. Front Aging Neurosci 12, 507140 (2020). 10.3389/fnagi.2020.507140

58 Abdellatif, M. et al. Fine-Tuning Cardiac Insulin-Like Growth Factor 1 Receptor Signaling to Promote Health and Longevity. Circulation 145, 1853–1866 (2022). doi:10.1161/CIRCULATIONAHA.122.059863

59 Jiao, Q., Takeshima, H., Ishikawa, Y. & Minamisawa, S. Sarcalumenin plays a critical role in age-related cardiac dysfunction due to decreases in SERCA2a expression and activity. Cell calcium 51, 31–39 (2012). 10.1016/j.ceca.2011.10.003

60 Li, S. et al. Cardiomyocytes disrupt pyrimidine biosynthesis in nonmyocytes to regulate heart repair. The Journal of clinical investigation 132 (2022). 10.1172/jci149711

61 Cai, W. et al. Alox15/15-HpETE Aggravates Myocardial Ischemia-Reperfusion Injury by Promoting Cardiomyocyte Ferroptosis. Circulation 147, 1444–1460 (2023). doi:10.1161/CIRCULATIONAHA.122.060257

62 Fan, Y. Z. et al. Vitronectin promotes TTR deposition and mitochondrial damage in ATTR-CA mice. European heart journal 44 (2023). 10.1093/eurheartj/ehad655.3191

63 Camacho-Pereira, J. et al. CD38 Dictates Age-Related NAD Decline and Mitochondrial Dysfunction through an SIRT3-Dependent Mechanism. Cell metabolism 23, 1127–1139 (2016). 10.1016/j.cmet.2016.05.006

64 Gardell, S. J. et al. Boosting NAD+ with a small molecule that activates NAMPT. Nature communications 10, 3241 (2019). 10.1038/s41467-019-11078-z

65 Luan, C. et al. Chloride intracellular channel CLIC3 mediates fibroblast cellular senescence by interacting with ERK7. Communications Biology 8, 51 (2025). 10.1038/s42003-025-07482-5

66 DeBerge, M. et al. Macrophage AXL receptor tyrosine kinase inflames the heart after reperfused myocardial infarction. The Journal of clinical investigation 131 (2021). 10.1172/jci139576

67 Singh, B. N. et al. A conserved HH-Gli1-Mycn network regulates heart regeneration from newt to human. Nature communications 9, 4237 (2018). 10.1038/s41467-018-06617-z

68 Waldron, C. J. et al. The HH-GLI2-CKS1B network regulates the proliferation-to-maturation transition of cardiomyocytes. Stem cells translational medicine 13, 678–692 (2024). 10.1093/stcltm/szae032

69 Luan, Y. et al. Cardiac cell senescence: molecular mechanisms, key proteins and therapeutic targets. Cell Death Discov 10, 78 (2024). 10.1038/s41420-023-01792-5

70 Shao, X. et al. CellTalkDB: a manually curated database of ligand–receptor interactions in humans and mice. Briefings in Bioinformatics 22 (2020). 10.1093/bib/bbaa269

71 Wilkins, J. T. & Rohatgi, A. Resolution of apolipoprotein A1 and A2 proteoforms: their cardiometabolic correlates and implications for future research. Current opinion in lipidology 33, 264–269 (2022). 10.1097/mol.0000000000000840

72 Happonen, K. E., Burrola, P. G. & Lemke, G. Regulation of brain endothelial cell physiology by the TAM receptor tyrosine kinase Mer. Commun Biol 6, 916 (2023). 10.1038/s42003-023-05287-y

73 dos Santos, L., et al. Circulating Dipeptidyl Peptidase IV Activity Correlates With Cardiac Dysfunction in Human and Experimental Heart Failure. Circulation: Heart Failure 6, 1029–1038 (2013). doi:10.1161/CIRCHEARTFAILURE.112.000057

74 Soare, A. et al. Dipeptidylpeptidase 4 as a Marker of Activated Fibroblasts and a Potential Target for the Treatment of Fibrosis in Systemic Sclerosis. Arthritis Rheumatol 72, 137–149 (2020). 10.1002/art.41058

75 Epstein, J. A., Aghajanian, H. & Singh, M. K. Semaphorin signaling in cardiovascular development. Cell metabolism 21, 163–173 (2015). 10.1016/j.cmet.2014.12.015

76 Zhou, B. et al. Adult mouse epicardium modulates myocardial injury by secreting paracrine factors. The Journal of clinical investigation 121, 1894–1904 (2011). 10.1172/jci45529

77 Molenaar, B. et al. Single-cell transcriptomics following ischemic injury identifies a role for B2M in cardiac repair. Commun Biol 4, 146 (2021). 10.1038/s42003-020-01636-3

78 Santos, M. et al. Defective iron homeostasis in beta 2-microglobulin knockout mice recapitulates hereditary hemochromatosis in man. J Exp Med 184, 1975–1985 (1996). 10.1084/jem.184.5.1975

79 Santos, M. M. et al. Iron overload and heart fibrosis in mice deficient for both beta2-microglobulin and Rag1. The American journal of pathology 157, 1883–1892 (2000). 10.1016/s0002-9440(10)64827-4

80 Liebermeister, W. et al. Visual account of protein investment in cellular functions. Proceedings of the National Academy of Sciences of the United States of America 111, 8488–8493 (2014). 10.1073/pnas.1314810111

81 Otto, A. et al. Systems-wide temporal proteomic profiling in glucose-starved Bacillus subtilis. Nature communications 1, 137 (2010). 10.1038/ncomms1137

82 Schäfer, S. et al. CD8(+) T Cells Drive Plaque Smooth Muscle Cell Dedifferentiation in Experimental Atherosclerosis. Arterioscler Thromb Vasc Biol 44, 1852–1872 (2024). 10.1161/atvbaha.123.320084

83 Smith, N. et al. Overlapping expression of Runx1(Cbfa2) and Runx2(Cbfa1) transcription factors supports cooperative induction of skeletal development. Journal of cellular physiology 203, 133–143 (2005). 10.1002/jcp.20210

84 Winslow, M. M. et al. Calcineurin/NFAT signaling in osteoblasts regulates bone mass. Dev Cell 10, 771–782 (2006). 10.1016/j.devcel.2006.04.006

85 Xu, Q. et al. Cellular communication network factor 2 regulates smooth muscle cell transdifferentiation and lipid accumulation in atherosclerosis. Cardiovascular research 120, 2191–2207 (2024). 10.1093/cvr/cvae215

86 Chakraborty, R. et al. Targeting smooth muscle cell phenotypic switching in vascular disease. JVS Vasc Sci 2, 79–94 (2021). 10.1016/j.jvssci.2021.04.001

87 Cappello, P. et al. CCL16/LEC powerfully triggers effector and antigen-presenting functions of macrophages and enhances T cell cytotoxicity. Journal of leukocyte biology 75, 135–142 (2004). 10.1189/jlb.0403146

88 Hauser, M. A. & Legler, D. F. Common and biased signaling pathways of the chemokine receptor CCR7 elicited by its ligands CCL19 and CCL21 in leukocytes. Journal of leukocyte biology 99, 869–882 (2016). 10.1189/jlb.2MR0815-380R

89 Mor, A. et al. Blockade of CCL24 with a monoclonal antibody ameliorates experimental dermal and pulmonary fibrosis. Ann Rheum Dis 78, 1260–1268 (2019). 10.1136/annrheumdis-2019-215119

90 Pulanco, M. C. et al. Complement Protein C1q Enhances Macrophage Foam Cell Survival and Efferocytosis. J Immunol 198, 472–480 (2017). 10.4049/jimmunol.1601445

91 Roy, A., Saqib, U., Wary, K. & Baig, M. S. Macrophage neuronal nitric oxide synthase (NOS1) controls the inflammatory response and foam cell formation in atherosclerosis. Int Immunopharmacol 83, 106382 (2020). 10.1016/j.intimp.2020.106382

92 Olshansky, S. J., Willcox, B. J., Demetrius, L. & Beltrán-Sánchez, H. Implausibility of radical life extension in humans in the twenty-first century. Nat Aging 4, 1635–1642 (2024). 10.1038/s43587-024-00702-3

93 Klösener, L. et al. Functional Cardiovascular Characterization of the Common Marmoset (Callithrix jacchus). Biology (Basel) 12 (2023). 10.3390/biology12081123

94 Luxán, G. & Dimmeler, S. The vasculature: a therapeutic target in heart failure? Cardiovascular research 118, 53–64 (2022). 10.1093/cvr/cvab047

95 Tamiato, A. et al. Age-Dependent RGS5 Loss in Pericytes Induces Cardiac Dysfunction and Fibrosis. Circulation research 134, 1240–1255 (2024). 10.1161/circresaha.123.324183

96 Zhang, F. et al. TGF-β induces M2-like macrophage polarization via SNAIL-mediated suppression of a pro-inflammatory phenotype. Oncotarget 7, 52294–52306 (2016). 10.18632/oncotarget.10561

97 Dadson, K. et al. Adiponectin mediated APPL1-AMPK signaling induces cell migration, MMP activation, and collagen remodeling in cardiac fibroblasts. Journal of cellular biochemistry 115, 785–793 (2014). 10.1002/jcb.24722

98 Menges, L. et al. Mind the gap (junction): cGMP induced by nitric oxide in cardiac myocytes originates from cardiac fibroblasts. British journal of pharmacology 176, 4696–4707 (2019). 10.1111/bph.14835

99 Menges, L. et al. It takes two to tango: cardiac fibroblast-derived NO-induced cGMP enters cardiac myocytes and increases cAMP by inhibiting PDE3. Commun Biol 6, 504 (2023). 10.1038/s42003-023-04880-5

100 Braunwald, E. Heart failure. JACC Heart Fail 1, 1–20 (2013). 10.1016/j.jchf.2012.10.002

101 Schimmel, K. et al. Natural Compound Library Screening Identifies New Molecules for the Treatment of Cardiac Fibrosis and Diastolic Dysfunction. Circulation 141, 751–767 (2020). 10.1161/circulationaha.119.042559

102 Iacobellis, G. Epicardial adipose tissue in contemporary cardiology. Nature reviews. Cardiology 19, 593–606 (2022). 10.1038/s41569-022-00679-9

103 Romero-Becera, R., Santamans, A. M., Arcones, A. C. & Sabio, G. From Beats to Metabolism: the Heart at the Core of Interorgan Metabolic Cross Talk. *Physiology (Bethesda*, Md.) 39, 98–125 (2024). 10.1152/physiol.00018.2023

104 Lopez, M. et al. Endothelial deletion of the cytochrome P450 reductase leads to cardiac remodelling. Frontiers in physiology 13, 1056369 (2022). 10.3389/fphys.2022.1056369

105 Klems, A. et al. The GEF Trio controls endothelial cell size and arterial remodeling downstream of Vegf signaling in both zebrafish and cell models. Nature communications 11, 5319 (2020). 10.1038/s41467-020-19008-0

106 Goettsch, C. et al. Nuclear factor of activated T cells mediates oxidised LDL-induced calcification of vascular smooth muscle cells. Diabetologia 54, 2690–2701 (2011). 10.1007/s00125-011-2219-0

107 Horstmann, H. et al. Cross-species single-cell RNA sequencing reveals divergent phenotypes and activation states of adaptive immunity in human carotid and experimental murine atherosclerosis. Cardiovascular research (2024). 10.1093/cvr/cvae154

108 Zhang, W. et al. A single-cell transcriptomic landscape of primate arterial aging. Nature communications 11, 2202 (2020). 10.1038/s41467-020-15997-0

109 Hinkel, R. et al. MRTF-A controls vessel growth and maturation by increasing the expression of CCN1 and CCN2. Nat Commun 5, 3970 (2014). 10.1038/ncomms4970

110 Paredes, F. et al. Metabolic regulation of the proteasome under hypoxia by Poldip2 controls fibrotic signaling in vascular smooth muscle cells. Free radical biology & medicine 195, 283–297 (2023). 10.1016/j.freeradbiomed.2022.12.098

111 Ismaeel, A. et al. Endothelial cell-derived pro-fibrotic factors increase TGF-β1 expression by smooth muscle cells in response to cycles of hypoxia-hyperoxia. Biochimica et biophysica acta. Molecular basis of disease 1868, 166278 (2022). 10.1016/j.bbadis.2021.166278

112 Raju, S. et al. Multiomics unveils extracellular vesicle-driven mechanisms of endothelial communication in human carotid atherosclerosis. bioRxiv (2024). 10.1101/2024.06.21.599781

113 He, X., Zeng, H. & Chen, J. X. Emerging role of SIRT3 in endothelial metabolism, angiogenesis, and cardiovascular disease. Journal of cellular physiology 234, 2252–2265 (2019). 10.1002/jcp.27200

114 Ungvari, Z., Tarantini, S., Sorond, F., Merkely, B. & Csiszar, A. Mechanisms of Vascular Aging, A Geroscience Perspective: JACC Focus Seminar. Journal of the American College of Cardiology 75, 931–941 (2020). 10.1016/j.jacc.2019.11.061

115 Kwak, H. B. Aging, exercise, and extracellular matrix in the heart. J Exerc Rehabil 9, 338–347 (2013). 10.12965/jer.130049

116 Franceschi, C. & Campisi, J. Chronic inflammation (inflammaging) and its potential contribution to age-associated diseases. J Gerontol A Biol Sci Med Sci 69 **Suppl 1**, S4–9 (2014). 10.1093/gerona/glu057

117 Dugan, B., Conway, J. & Duggal, N. A. Inflammaging as a target for healthy ageing. Age and ageing 52 (2023). 10.1093/ageing/afac328

118 Sanada, F. et al. Source of Chronic Inflammation in Aging. Frontiers in cardiovascular medicine 5, 12 (2018). 10.3389/fcvm.2018.00012

119 Lima, V. L., Sena, V. L., Stewart, B., Owen, J. S. & Dolphin, P. J. An evaluation of the marmoset Callithrix jacchus (sagüi) as an experimental model for the dyslipoproteinemia of human Schistosomiasis mansoni. Biochimica et biophysica acta 1393, 235–243 (1998). 10.1016/s0005-2760(98)00076-9

120 Crook, D., Weisgraber, K. H., Boyles, J. K. & Mahley, R. W. Isolation and characterization of plasma lipoproteins of common marmoset monkey. Comparison of effects of control and atherogenic diets. Arteriosclerosis 10, 633–647 (1990). 10.1161/01.atv.10.4.633

121 Ross, C. N. et al. Cross-sectional comparison of health-span phenotypes in young versus geriatric marmosets. Am J Primatol 81, e22952 (2019). 10.1002/ajp.22952

122 Plazyo, O., Hao, W. & Jin, J. P. The Absence of Calponin 2 in Rabbits Suggests Caution in Choosing Animal Models. Frontiers in bioengineering and biotechnology 8, 42 (2020). 10.3389/fbioe.2020.00042

123 Knowles, J. W. & Maeda, N. Genetic modifiers of atherosclerosis in mice. Arteriosclerosis, thrombosis, and vascular biology 20, 2336–2345 (2000). 10.1161/01.atv.20.11.2336

124 Marotti, K. R. et al. Severe atherosclerosis in transgenic mice expressing simian cholesteryl ester transfer protein. Nature 364, 73–75 (1993). 10.1038/364073a0

125 Mietsch, M. et al. Blood pressure as prognostic marker for body condition, cardiovascular, and metabolic diseases in the common marmoset (Callithrix jacchus). J Med Primatol 45, 126–138 (2016). 10.1111/jmp.12215

126 Mattison, J. A. & Vaughan, K. L. An overview of nonhuman primates in aging research. Experimental gerontology 94, 41–45 (2017). 10.1016/j.exger.2016.12.005

127 Teo, K. K. & Rafiq, T. Cardiovascular Risk Factors and Prevention: A Perspective From Developing Countries. The Canadian journal of cardiology 37, 733–743 (2021). 10.1016/j.cjca.2021.02.009

128 Abbott, D. H., Barnett, D. K., Colman, R. J., Yamamoto, M. E. & Schultz-Darken, N. J. Aspects of common marmoset basic biology and life history important for biomedical research. Comp Med 53, 339–350 (2003).

129 Cui, M. & Olson, E. N. Protocol for Single-Nucleus Transcriptomics of Diploid and Tetraploid Cardiomyocytes in Murine Hearts. STAR Protoc 1, 100049 (2020). 10.1016/j.xpro.2020.100049

130 Jin, S. et al. Inference and analysis of cell-cell communication using CellChat. Nature communications 12, 1088 (2021). 10.1038/s41467-021-21246-9

131 Bleckwedel F, G. G., Zelarayan LC. An optimized protocol for the enrichment of small vesicles from murine and non-human primate heart tissue. Trillium Extracellular Vesicles 4 (2022). 10.47184/tev.2022.01.03

132 Görgens, A. et al. Identification of storage conditions stabilizing extracellular vesicles preparations. J Extracell Vesicles 11, e12238 (2022). 10.1002/jev2.12238

133 Théry, C., Amigorena, S., Raposo, G. & Clayton, A. Isolation and characterization of exosomes from cell culture supernatants and biological fluids. Curr Protoc Cell Biol Chapter 3, Unit 3.22 (2006). 10.1002/0471143030.cb0322s30

134 Atanassov, I. & Urlaub, H. Increased proteome coverage by combining PAGE and peptide isoelectric focusing: comparative study of gel-based separation approaches. Proteomics 13, 2947–2955 (2013). 10.1002/pmic.201300035

135 Dobin, A. et al. STAR: ultrafast universal RNA-seq aligner. Bioinformatics 29, 15–21 (2013). 10.1093/bioinformatics/bts635

136 Liao, Y., Smyth, G. K. & Shi, W. featureCounts: an efficient general purpose program for assigning sequence reads to genomic features. Bioinformatics 30, 923–930 (2014). 10.1093/bioinformatics/btt656

137 Love, M. I., Huber, W. & Anders, S. Moderated estimation of fold change and dispersion for RNA-seq data with DESeq2. Genome Biol 15, 550 (2014). 10.1186/s13059-014-0550-8

138 Edgar, R., Domrachev, M. & Lash, A. E. Gene Expression Omnibus: NCBI gene expression and hybridization array data repository. Nucleic Acids Res 30, 207–210 (2002). 10.1093/nar/30.1.207

139 Perez-Riverol, Y. et al. The PRIDE database at 20 years: 2025 update. Nucleic Acids Research 53, D543–D553 (2024). 10.1093/nar/gkae1011

