## supplementary figures for "Natural aging drives a subclinical cardiovascular phenotype in a non-human primate"

**Supplementary Figure 1: Body and heart weights and blood parameters of young-adult and aged animals.**

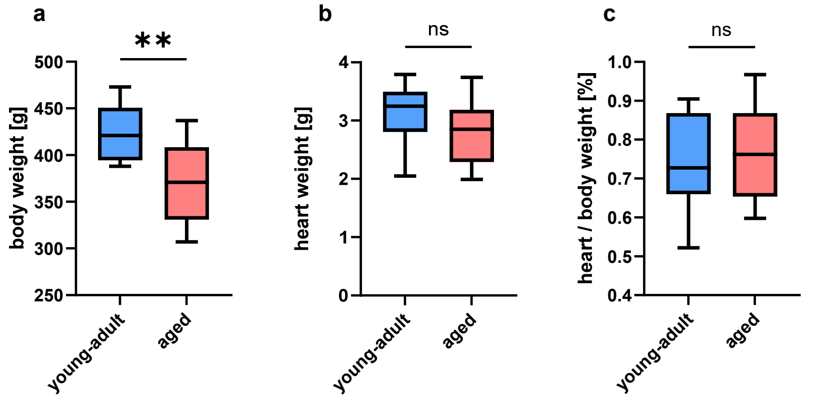


**Measurements at initial examination:**

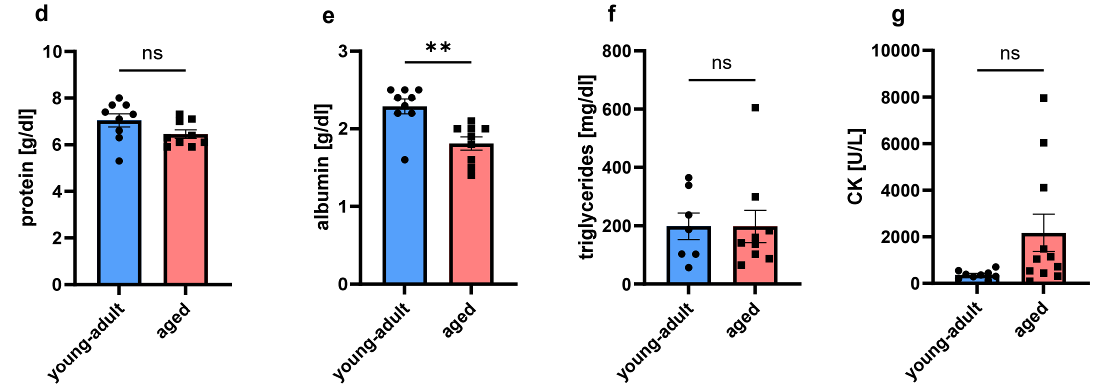


**Final measurements previous to necropsy:**

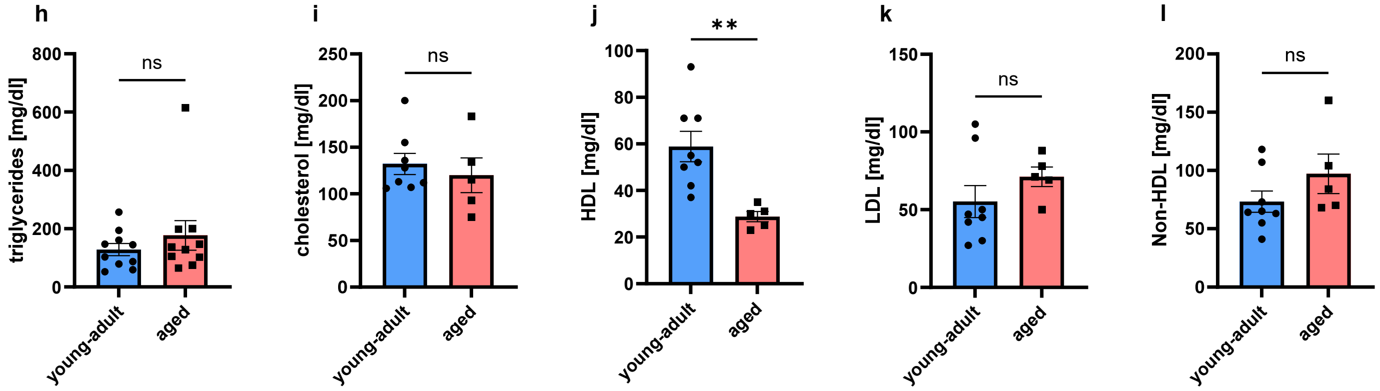


**a**, Body weight, **b**, heart weight and **c**, heart to body weight relation (n = 9; whiskers = min-max values). **d**–**g** Results for blood parameters at initial examination (n = 9). **h**–**i** Blood markers of lipid metabolismat (young-adult n = 8, aged n = 5) at the timepoint of necropsy: triglycerides (**h**); total cholesterol (**i**); high density lipoprotein (HD) (**j**); low density lipoprotein (LDL) (**k**); and non-HDL (**l**). Bar graphs in **d**–**l** represent means ±SEM. Statistical significance was assessed by unpaired two-tailed Student's *t*-tests (**P* < 0.05, ***P* < 0.01, ns: not significant).

**Supplementary Figure 2: Exemplary images of intra-cardiac pressure-volume (PV) measurements and histology.**

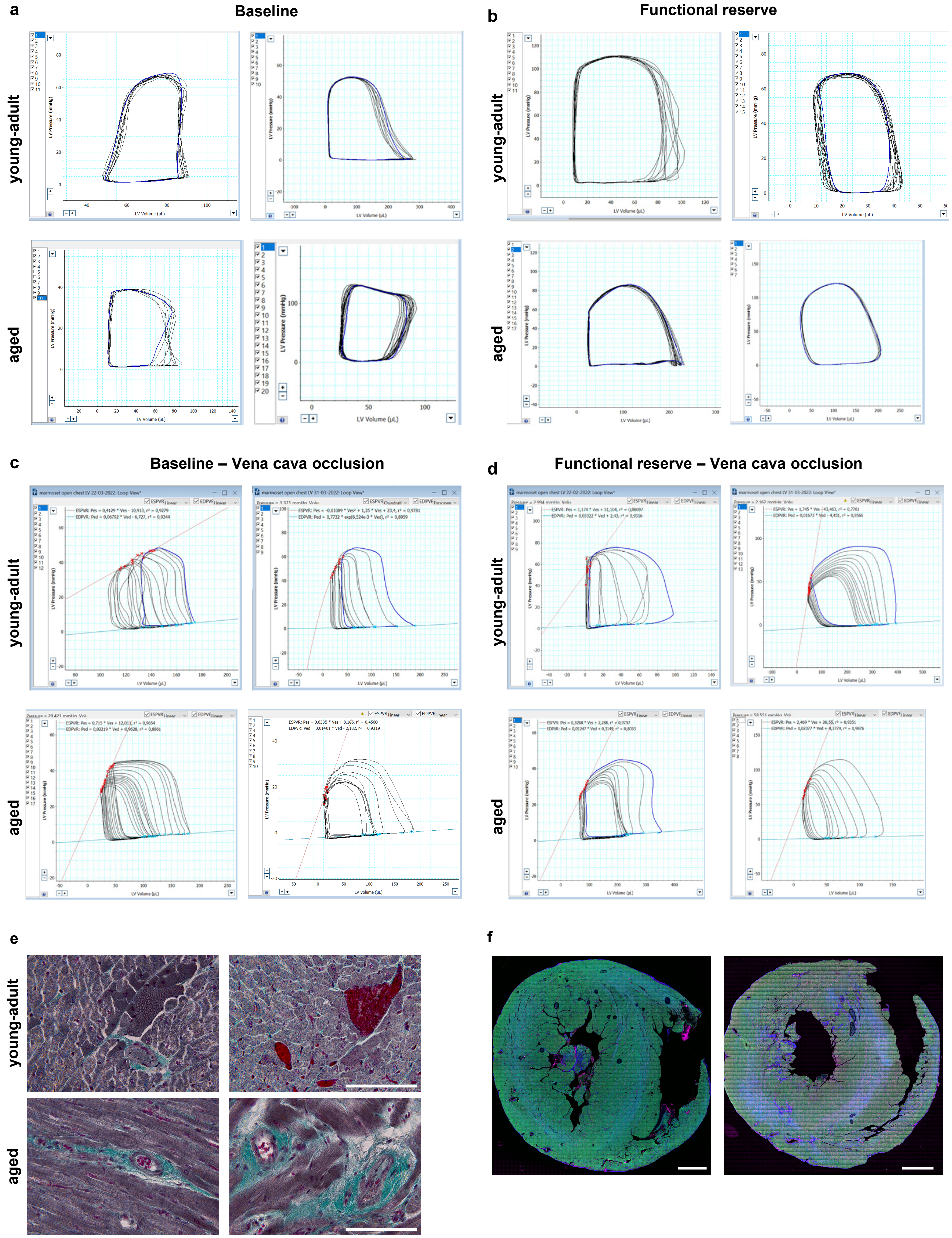


**a**,**b**, Representative images of PV measurements at baseline (**a**) and functional reserve (**b**). **c**,**d** Representative images of vena cava occlusion measurements to assess load independent heart function at baseline (**c**) and functional reserve (**d**). **e**, Representative images of Masson Trichrome Staining for examination of perivascular fibrosis. Scale bar = 100 µm. **f**, Representative label-free three-photon microscopy pictures used for microvasculature analysis. Scale bar = 2 mm.

**Supplementary Figure 3: Investigation of atherosclerosis.**

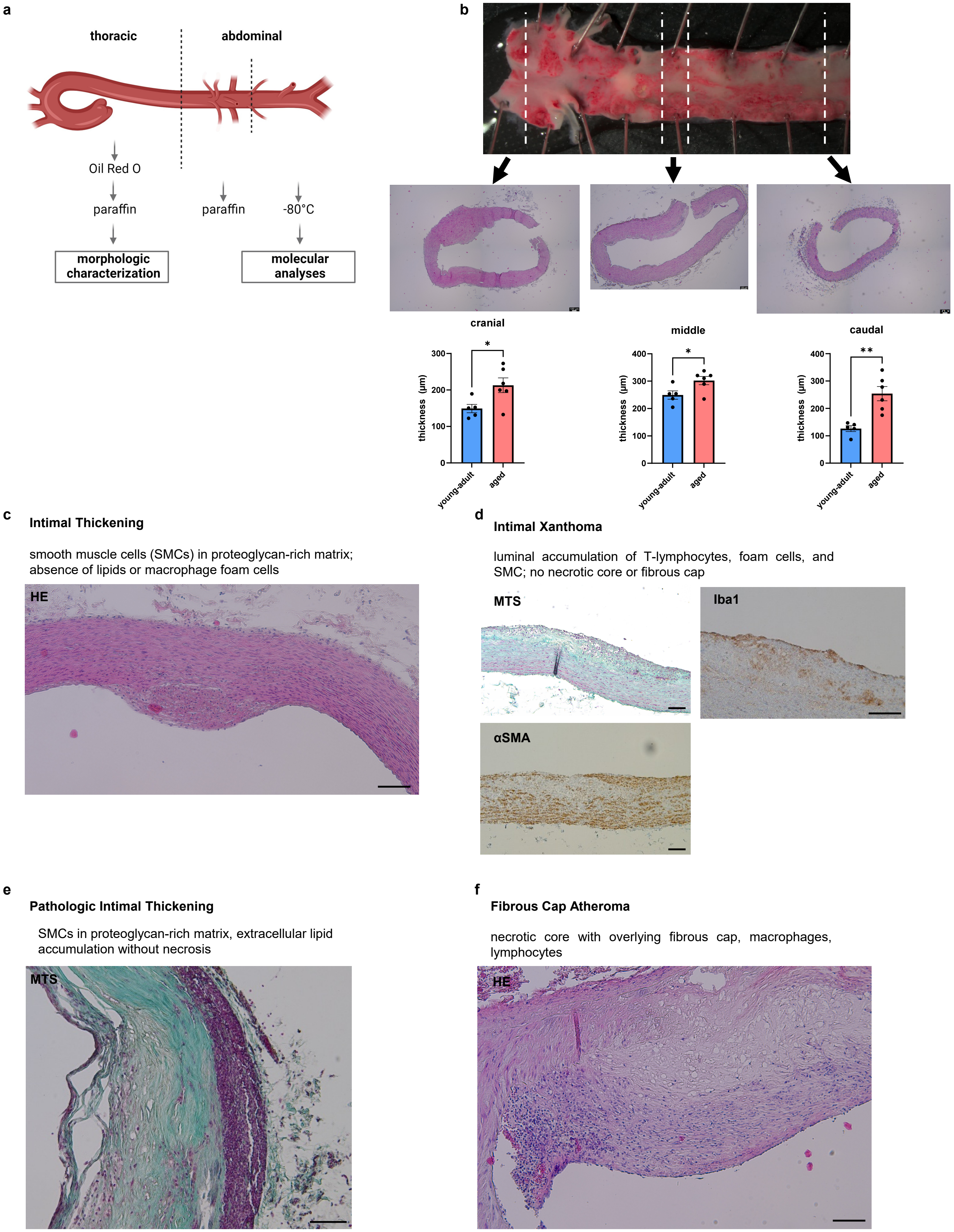


**a**, Sampling scheme of aortas for further analyses. **b**, Analysis of three localisations within the thoracic aorta for wall thickness measurement and respective results for each location. **c-f**, Examples for plaque grading according to Virmani et al.2000. Scale bars = 100 µm.

**Supplementary Figure 4: Single-nuclei RNA-sequencing of young-adult or aged marmoset heart tissue.**

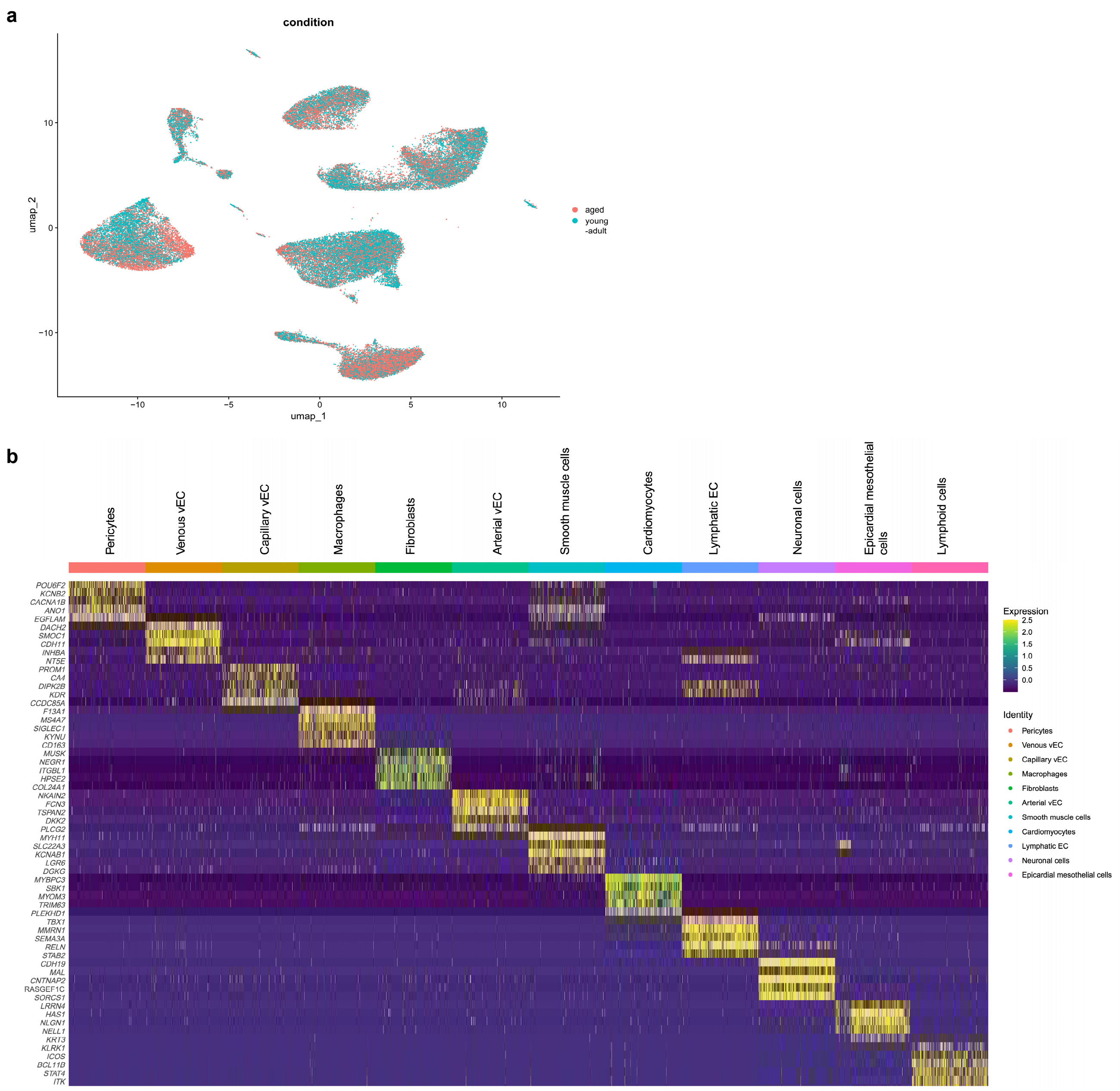


**a**, UMAP projection of snRNA-seq illustration of distinct spatial clustering of cells from young-adult and aged animals. **b**, Heatmap displaying the expression signature of the top 5 cell type-specific genes per cluster.

**Supplementary Figure 5: Differential gene expression and enrichment analysis of cardiac cell clusters from young-adult or aged animals by snRNA-seq.**

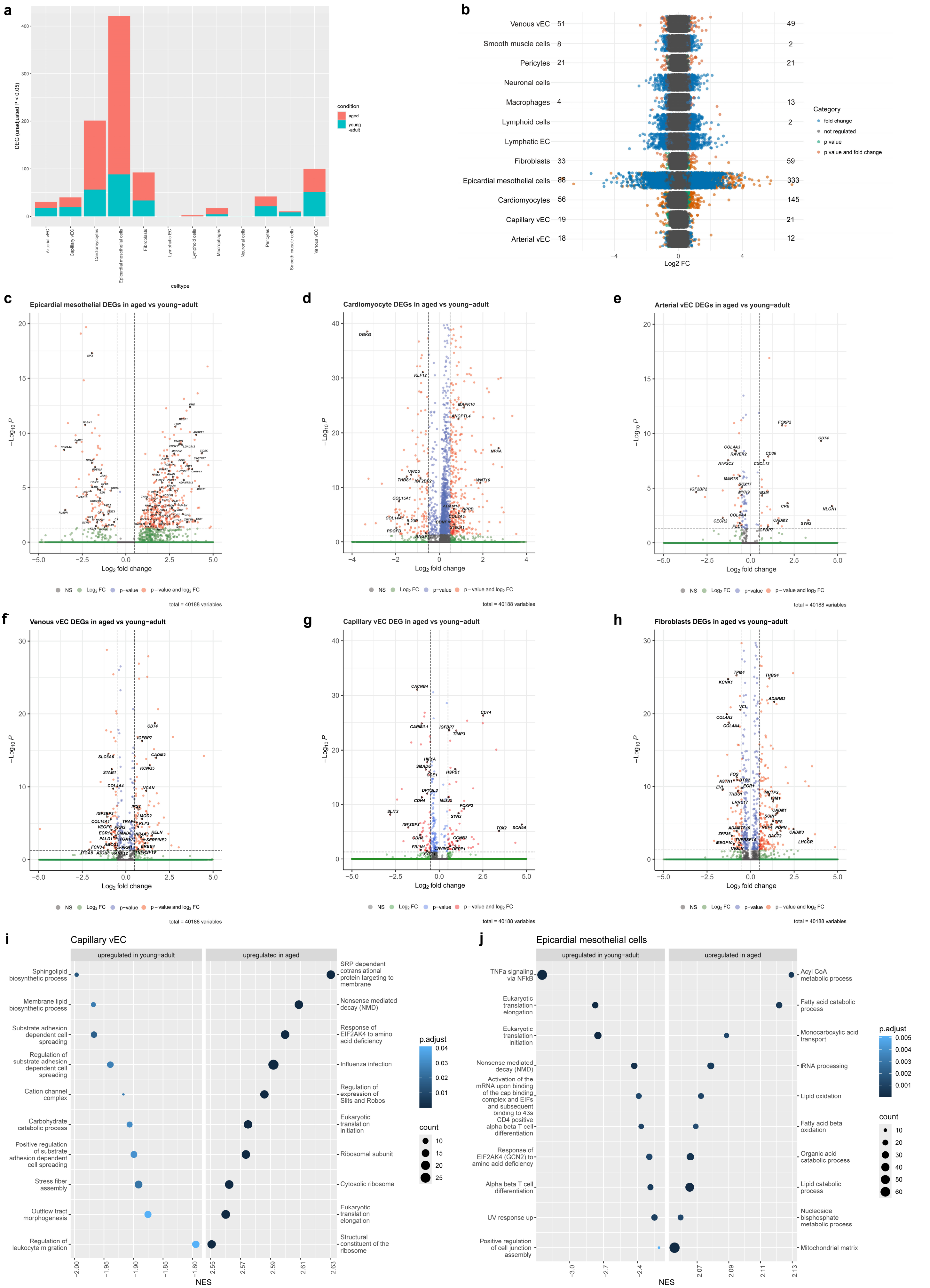


**a**,**b**, Stacked bar and scatter plots showing the count of differentially expressed genes in different cardiac clusters in young-adult or aged animals. **c**–**h**, Volcano plots showing differentially expressed genes in (**c**) epicardial mesothelial cells; (**d**) cardiomyocytes; (**e**) arterial vascular endothelial cells (vEC); (**f**) venous vEC; (**g**) capillary vEC and (**h**) fibroblasts in young-adult or aged hearts. **i**,**j**, Gene set enrichment analysis (GSEA) of differentially expressed genes in capillar vEC and epicardlal mesothelial cells in young-adult or aged hearts.

**Supplementary Figure 7: Cardiac inter-cellular signaling by CellChat analysis in young-adult or aged animals.**

**a**

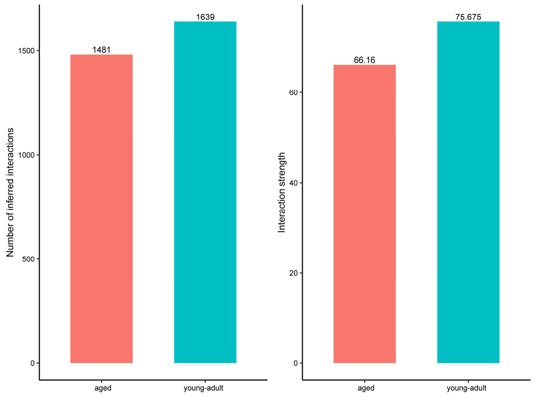


**b**

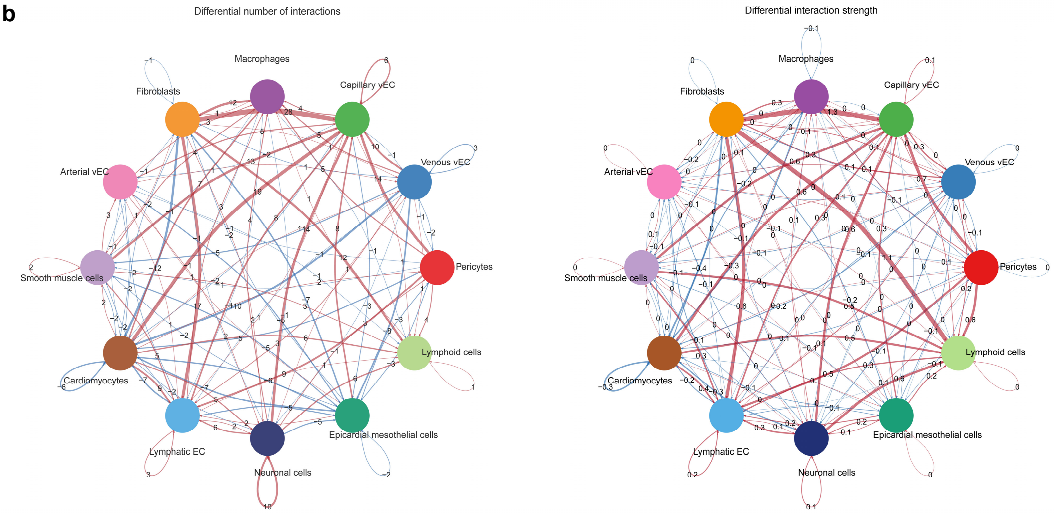


**a**, Bar plot comparing cardiac inter-cellular signal count and strength in young-adult or aged animals. **b**, Diiferential interaction circle plot of cardiac inter-cellular signal count and strength in young-adult vs. aged animals.

**Supplementary Figure 7: Proteomic analysis of cardiac extracellular vesicles (EVs) from young-adult or aged animals.**

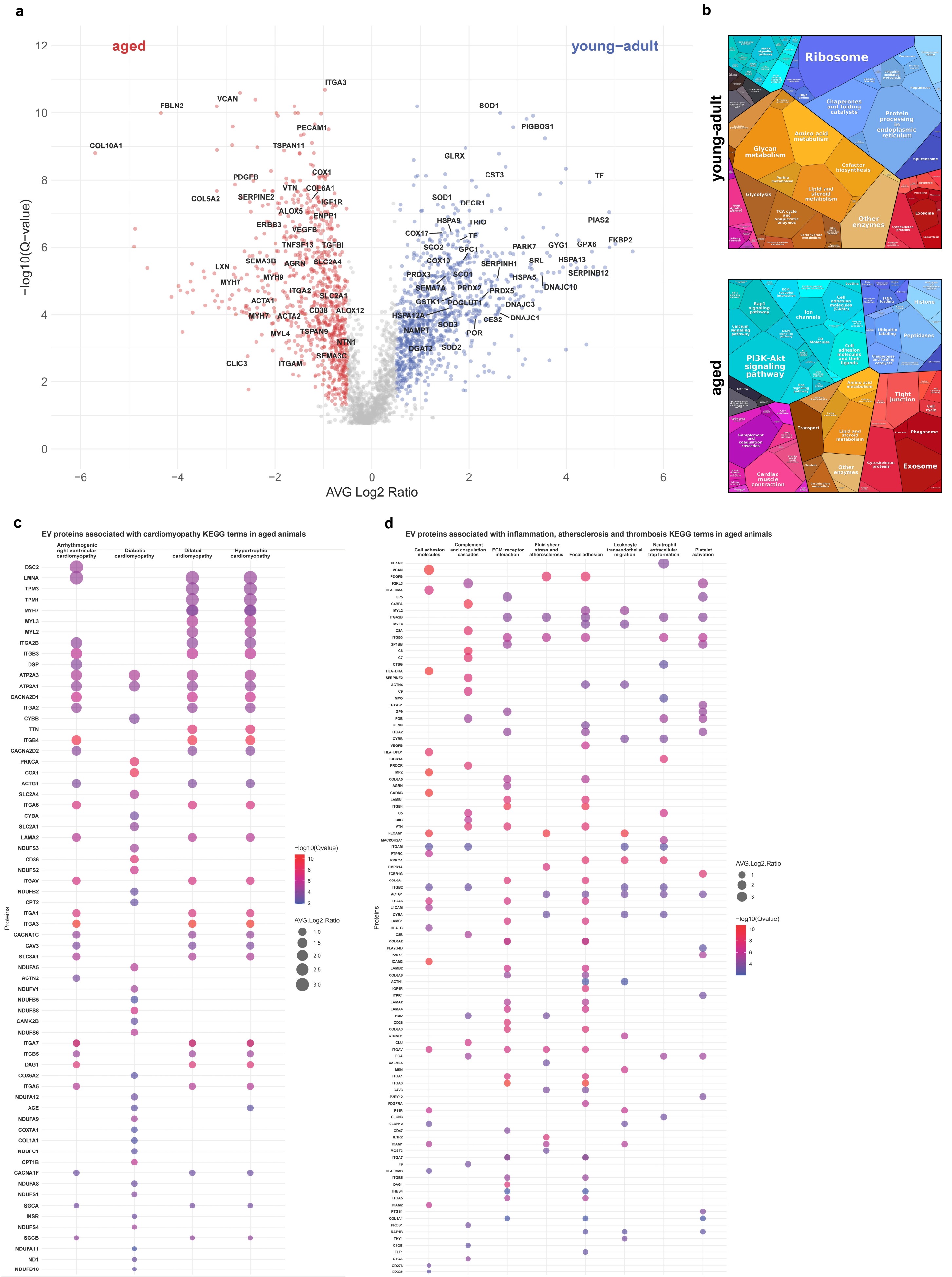


**a**, Volcano plot of differentially regulated proteins of cardiac EVs from young-adult or aged animals. **b**, Proteomap enrichment analysis of differentially expressed proteins in cardiac EVs from young-adult or aged animals. **c**,**d**, Dot plots of KEGG analysis of differentially upregulated proteins in aged cardiac EVs by proteomic analysis, showing proteins enriched for terms of four cardiomyopathies (arrhythmogenic, diabetic, dilated and/or hypertrophic cardiomyopathy) (**c**); and terms associated with inflammation, atherosclerosis and thrombosis (**d**).
